# An ultra high-density *Arabidopsis thaliana* crossover map that refines the influences of structural variation and epigenetic features

**DOI:** 10.1101/665083

**Authors:** Beth A. Rowan, Darren Heavens, Tatiana R. Feuerborn, Andrew J. Tock, Ian R. Henderson, Detlef Weigel

## Abstract

Many environmental, genetic, and epigenetic factors are known to affect the frequency and positioning of meiotic crossovers (COs). Suppression of COs by large, cytologically visible inversions and translocations has long been recognized, but relatively little is known about how smaller structural variants (SVs) affect COs. To examine fine-scale determinants of the CO landscape, including SVs, we used a rapid, cost-effective method for high-throughput sequencing to generate a precise map of over 17,000 COs between the Col-0 and Ler accessions of *Arabidopsis thaliana*. COs were generally suppressed in regions with SVs, but this effect did not depend on the size of the variant region, and was only marginally affected by the variant type. CO suppression did not extend far beyond the SV borders, and CO rates were slightly elevated in the flanking regions. Disease resistance gene clusters, which often exist as SVs, exhibited high CO rates at some loci, but there was a tendency toward depressed CO rates at loci where large structural differences exist between the two parents. Our high-density map also revealed in fine detail how CO positioning relates to genetic (DNA motifs) and epigenetic (chromatin structure) features of the genome. We conclude that suppression of COs occurs over a narrow region spanning large and small-scale SVs, representing influence on the CO landscape in addition to sequence and epigenetic variation along chromosomes.

## Introduction

Sexual reproduction generates genetic diversity from standing variation because meiotic crossovers (COs) between the maternal and paternal chromosomes provide new combinations of alleles that can be transmitted to the next generation (Barton and Charlesworth 1998). This novel variation can contribute to phenotypes upon which selection can act (Burt 2000), bring together beneficial mutations that arose independently (Muller 1932; Fisher 1999), and it can break linkage between beneficial and deleterious mutations (Peck 1994; Gray and Goddard 2012). Therefore, sexual reproduction, through the action of COs, provides a mechanism for accelerating adaptation. COs ensure proper chromosome segregation (Hall 1972) and are one result of several alternative outcomes of the repair of programmed DNA double-strand breaks (DSBs) that occur during meiosis that also include non-crossover and inter-sister repair (Szostak *et al*. 1983; Keeney *et al*. 1997; Keeney 2001). Therefore, a deeper understanding of the various influences affecting the repair of meiotic DSBs, especially those that favor the generation of COs, will strengthen our knowledge of processes that shape adaptive genetic variation.

One important factor that influences the positioning of COs along chromosomes, collinearity, has been known for almost a century. Alfred Sturtevant first proposed that a rearrangement of chromosome structure, such as an inverted segment, would suppress CO formation in the heterozygous state (Sturtevant 1921). He later confirmed that a region of chromosome III in *Drosophila* that exhibited CO suppression in the heterozygous state did indeed contain an inversion (Sturtevant 1926). Such inverted regions can encompass many genes, with the consequence that alleles in the inverted segment are passed down as a single non-recombining locus. When genes that are linked in such a way confer a particularly advantageous trait as a single unit, they form a supergene (Schwander *et al*. 2014; Thompson and Jiggins 2014; Charlesworth 2016) — a concept that had its origins in Dobzhansky’s work and was formalized by Mather (Mather 1950). A notable example from the animal kingdom comes from the ruff (*Philomachus pugnax*), where an inversion has trapped 125 genes in a non-recombining haplotype, creating a supergene that governs both ornamental plumage and mating/social behaviors (Küpper *et al*. 2016). Another inversion-based supergene underlies wing pattern morphology and mimicry in butterflies (Joron *et al*. 2011). In plants, a chromosomal inversion is responsible for variation in life-history strategies and adaptation to temperature and precipitation in yellow monkeyflowers (Lowry and Willis 2010; Oneal *et al*. 2014). Inversions are not the only type of chromosomal rearrangement resulting in CO suppression. Dobzhansky first observed reduced crossing over near interchromosomal translocations in *Drosophila* (Dobzhansky 1931), and this has been observed in other species (McKim *et al*. 1988; Herickhoff *et al*. 1993). Interestingly, some translocations can enhance recombination in intervals flanking the breakpoint (Sybenga 1970).

Smaller-scale structural variants (SVs) of different types have also been implicated in suppression of meiotic recombination. For example, a 70-kb transposition on *Arabidopsis thaliana* chromosome 3 showed extreme local suppression of recombination at the excision site in an F_2_ cross between accessions BG-5 and Kro-0 (Alhajturki *et al*. 2018). This type of rearrangement creates an insertion/deletion (indel) polymorphism at both the original locus and the new location. It is therefore expected that other types of insertions or deletions would also show local CO suppression. Indeed, transgene arrays on the order of dozens of kilobases induced local CO suppression in the nematode *Caenorhabditis elegans* (Hammarlund *et al*. 2005). In mouse, the co-occurence of a small insertion (824 bp) and a small deletion (703 bp) was associated with a local reduction in the crossover rate (Hsu *et al*. 2000).

Despite a large body of literature supporting a role for structural variation in determining CO positioning, there has been surprisingly little systematic investigation of whether the size or the type of variant has a differential impact on the local CO rate. In *Drosophila* interspecies crosses, suppression of meiotic COs can extend for more than 1 Mb outside the borders of inversions (Kulathinal *et al*. 2009; Stevison *et al*. 2011). However, it is unclear how far beyond the variant borders the effects of CO suppression can extend in other species or for other types of SVs. In this study, we developed a new fast and cost-effective protocol for preparation of Illumina genomic DNA sequencing libraries. With this method, we sequenced the genomes of nearly 2,000 F_2_ individuals from a cross between the *A. thaliana* accessions Col-0 and L*er*. After including previously published sequence data for additional F_2_ recombinants from this cross (Choi *et al*. 2016; Underwood *et al*. 2018), we generated a set of over 17,000 COs at a high resolution (median: 1,102 bp) and examined them in the context of the precise knowledge of SVs available from high-quality reference genome sequences for these two accessions. We then compared the relationship between COs and SVs to other known influences on recombination rate.

## Materials and Methods

### Plant growth and DNA extraction

Col-0 *qrt1-2 CEN3 420 (Melamed-Bessudo et al. 2005; Francis et al. 2007)* x L*er* F_2_ seeds (Ziolkowski *et al*. 2015) were stratified for 4-7 days at 4°C before sowing on soil in 40-pot trays. A portion of the seeds (~400) had been subjected to screening for a lack of COs in the *420* crossover reporter region on chromosome 3. Plants were cultivated in a greenhouse supplemented with additional light and covered with a plastic dome for the first week of growth to promote germination and establishment. Leaf samples (approximately 0.5-1 cm long) were taken from plants between 4-8 weeks of age, collected in polypropylene tubes in a 96-well rack, and frozen at −80°C until DNA extraction.

To extract DNA, the frozen leaf samples were ground using a TissueLyser (Qiagen Hilden, Germany) with steel beads. A volume of 500 μL of DNA Extraction Buffer (200 mM Tris-HCl, 25 mM EDTA,1% SDS, 80 μg/mL RNase A) was added to the frozen powder and the tubes were mixed by gentle inversion before incubation at 37°C for 1 hour. The tubes were centrifuged at 3,000 x g for 5 minutes (min) and a volume of 400 μL of the supernatant was transferred to each well of a 96-deep-well plate containing 130 uL of potassium acetate solution (5M potassium acetate and 7% Tween-20); this and the following steps were performed with the aid of a Tecan Freedom EVO 150 liquid handling robot (Tecan, Männedorf, Switzerland) fitted with an 8 channel LiHa, a 96-channel MCA, and a gripper. Plates were covered with an adhesive seal and mixed gently by inversion before incubation for 10 min on ice and centrifugation for 5 min at 3,000 x g. A volume of 400 μL of the supernatant was transferred to a fresh 96-deep-well plate containing 400 uL of SeraPure solid-phase reverse immobilization (SPRI) beads (Rowan *et al*. 2017) and mixed several times by pipetting using a Tecan liquid-handling robot before incubation for 5 min at room temperature to allow the DNA to bind to the beads. The plate was placed on a magnetic plate stand, and the supernatant was removed and discarded after all of the beads had been drawn to the magnet (~5 min). The beads were washed three times with 900 μL of 80% EtOH. After the last wash, all traces of EtOH were removed and the beads were left to dry at room temperature for 5 min. The DNA was eluted from the beads in 100 μL of 10 mM Tris-HCl, pH 8, and mixed for 1 min using a plate vortex mixer and incubated overnight at 4°C. The plates were placed on a magnet for 5 min until the samples were clear of beads and the DNA solutions were transferred to a 96-well PCR plate and stored at −20°C until use in library prep.

### Library preparation and sequencing using Nextera LITE

DNA samples were first quantified using the Quant-It kit (Thermo Fisher Scientific, Waltham, MA) with a Tecan (Männedorf, Switzerland) Infinite M200 Pro fluorescence microplate reader (Tecan, (Männedorf, Switzerland) and the Tecan Magellan analysis software. The samples were normalized to 2 ng/μL using a Tecan liquid-handling robot. After normalization, a subset of samples from each plate was quantified using a Qubit 2.0 fluorometer with the dsDNA High Sensitivity assay kit in order to verify the concentration and assess the variability among samples. If the concentrations were on average 2 ng/μL with less than a two-fold variation among samples, then all samples in a plate were diluted uniformly to 0.25 - 0.5 ng/μL. Otherwise, the normalization was repeated. Once the samples were at 0.25 - 0.5 ng/μL, then 2 μL of DNA per sample were transferred to a fresh plate and mixed with 2.1 μL water, 0.82 μL TD buffer and 0.08 μL TD enzyme from the Illumina Nextera 24-sample kit (Illumina, Inc., San Diego, CA, USA). Plates were sealed with a Microseal B adhesive plate seal (Bio-Rad, Hercules, CA, USA) and incubated for 10 min at 55°C. After allowing the plates to cool to room temperature, the tagmented DNA was amplified by PCR using the KAPA2G Robust PCR kit (Sigma-Aldrich, St. Louis, MO, USA) using the GC buffer along with 0.2 μM of each custom P5 and P7 indexing primers (File S1) using the following cycling conditions: 72°C for 3 min, 95°C for 1 min, 14 cycles of 95°C for 10 s, 65°C for 20 s, and 72°C for 3 min.

Following PCR, a 5-μL sample of each library was mixed with 15 µL of 10 mM Tris-HCl, pH 8, and 5 µL of 6x loading dye and analyzed by electrophoresis using a 2% agarose gel at 140 V for 15 min to evaluate library success and size distribution. Samples were then pooled by mixing 3 μL of each library into a single tube using a Tecan liquid-handling robot. Note that the indexing oligos in File S1 allow for 576 unique combinations of P5 and P7 indices, but additional oligos can be easily designed if higher levels of multiplexing are desired.

Size selection was performed on 100 μL of the 96-sample pool by adding 160 μL of Serapure SPRI beads (Rowan *et al*. 2017) that had been diluted to 0.6x strength with water and incubating for 5 min at room temperature before placing in a magnetic 1.5-mL tube stand for 5 min. A volume of 250 μL of the supernatant was transferred to a fresh tube and 700 μL of 80% EtOH were added to the tube containing the beads (bead fraction 1). A volume of 30 μL full-strength Serapure SPRI beads was added to the supernatant and incubated for 5 minutes at room temperature and then for 5 min in a magnetic 1.5-mL tube stand before transferring 280 μL of the supernatant to a fresh tube. A volume of 700 μL of 80% EtOH was added to the beads that remained after this transfer (bead fraction 2). A volume of 112 μL full-strength Serapure SPRI beads was added to the supernatant and incubated for 5 min at room temperature and for 5 min in a magnetic 1.5-mL tube stand before removing and discarding the supernatant. A volume of 700 μL of 80% EtOH was added to the beads that remained after this transfer (bead fraction 3) and the tubes were incubated for at least 30 s before removing the EtOH from all bead fractions and doing a second wash with 700 μL of 80% EtOH. After at least 30 s in the second EtOH wash, all EtOH was removed from the tubes. A flash spin was performed to collect any remaining EtOH at the bottom of the tube, which was removed using a pipet. The bead samples were left to dry at room temperature for 5 min or until no EtOH remained.

The libraries for each of the three bead fractions were eluted in 26 μL of 10 mM Tris-HCl and incubated in a standard tube rack for 5 min. The tubes were then transferred back to the magnetic rack and 25 µL of the eluted libraries for each fraction were transferred to a fresh tube after all of the beads had been drawn to the magnet. Each of the three size fractions was analyzed on a Bioanalyzer to determine the size ranges and select the appropriate fraction for Illumina sequencing (often bead fraction 3, which contained 300-500 bp fragments). Samples were sequenced to an estimated depth of 1-2x average whole-genome coverage on an Illumina HiSeq 3000 instrument.

### Sequence analysis and crossover localization

The raw reads were demultiplexed and evaluated for quality using MultiQC v. 0.9.dev0 (Ewels *et al*. 2016) before adapter trimming and quality filtering using Trimmomatic v. 0.36 (Bolger *et al*. 2014). The reads were then aligned to the *A. thaliana* Col-0 reference genome (TAIR10) with BWA (Li and Durbin 2009) v0.6.2 using the *bwa aln* algorithm with default parameters, but specifying a maximum of 10 mismatches. Demultiplexed raw reads for an additional 192 wild-type Col X Ler F_2_ samples were downloaded from http://www.ebi.ac.uk/arrayexpress/experiments/E-MTAB-4657/samples/ and for an additional 171 Col X Ler F_2_ samples from http://www.ebi.ac.uk/arrayexpress/experiments/E-MTAB-5476/samples/ and subjected to the same methods of quality filtering and alignment. Paired-end reads were aligned independently, then merged using *sampe* with the default parameters and specifying a maximum insert size of 500 bp.

SAMtools v.1.2 and BCFtools v.1.2 (Li *et al*. 2009) were used to obtain read count data for variant positions to generate the input files for the TIGER crossover analysis pipeline (Rowan *et al*. 2015). A set of 545,481 total variants including single nucleotide polymorphisms (SNPs) and small (1-3 bp) insertion-deletion polymorphisms (indels) based on assembly comparisons between Ler and Col was filtered to SNPs only to generate the “complete” marker file (519,525 positions). The intersection between this and a set of 291,973 high-quality Col/Ler SNPs based on short read data (kindly provided by Korbinian Schneeberger, Max Planck Institute for Plant Breeding Research) yielded a final set of 237,288 SNPs as the “corrected” marker file.

Using this set as the input for TIGER, breakpoints were predicted on each sample independently, with the biparental mode setting and default parameters (Rowan *et al*. 2015). Briefly, genotypes were called for individual high-quality marker positions with the TIGER Basecaller module, and the sequence of genotypes along the chromosomes was reconstructed for each individual sample with the TIGER Hidden Markov Model, using a sample-specific error probably model generated from a beta-binomial model, with which we inferred the underlying genotype probability based on the allele frequency distributions. Final recombination breakpoint resolution was refined by gathering information from additional markers (the “complete” marker set described above) around the breakpoints. Refined breakpoint data files for all individuals were concatenated into a single text file for downstream analysis in R. Individuals with less than 0.025x coverage, or with excessive breakpoints, or without breakpoints, or with extended marker blocks where the allele frequency was not 0, 0.5 or 1, were removed from further analysis.

For downstream analysis, a crossover (CO) “interval” was designated as the region in between the last marker of one genotype and the first marker of the new genotype. CO positions were estimated as the midpoint between the positions of these two markers. CO positions that appeared to be double COs less than 500 kb apart were removed as likely false positives.

We obtained the positions and annotations of Col/Ler SVs from a Ler genome assembly (Zapata *et al*. 2016). The positions and annotations for disease resistance genes (including nucleotide-binding-site-leucine-rich-repeat proteins (NLRs)) for Col were obtained from (Choi *et al*. 2016), and we performed a comparative analysis with Ler annotations (Korbinian Schneeberger, personal communication) to determine NLR copy number variation and SVs. The DNase I hypersensitive hotspot data for 7-day-old seedlings (Sullivan *et al*. 2014) were downloaded from http://plantregulome.org/. SPO11-1-oligo and micrococcal nuclease sequencing (MNase-seq) data were obtained from (Choi *et al*. 2016, 2018). Roger Deal and Marco Bajkic kindly provided raw Assay for Transposase Accessible Chromatin sequencing (ATAC-seq) data from (Maher *et al*. 2018) and the data generated from meristem tissue was used in our analyses.

Statistical analysis of CO intervals and CO positions were performed using the R software environment (R Core Team 2017). Centromeres were defined as Chr1: 13,700,000..15,900,000; Chr2: 2,450,000..5,500,000; Chr3: 11,300,000..14,300,000; Chr4: 1,800,000..5,150,000; and Chr5: 11,000,000..13,350,000. Overlaps between CO intervals and the regions of SVs, NLR loci, SPO11-1-oligo hotspots, and ATAC-seq sites were analyzed with regioneR (Gel *et al*. 2016). Additional plots were prepared using ggplot2 (http://ftp.auckland.ac.nz/software/CRAN/src/contrib/Descriptions/ggplot.html). GO enrichment in CO deserts was performed using PANTHER version 13.1 (http://pantherdb.org/). COs, structural variants were grouped into 10-kb fixed windows and compared to DNA methylation data (Yelina *et al*. 2015) for the same genomic windows. Motif enrichment analyses were performed with MEME (Bailey *et al*. 2009) after subsetting the CO data to 500 and 1000 randomly selected COs per chromosome in order to achieve a feasible runtime. Parameters for MEME were set to search for motifs of lengths 2 to 10 bp with zero or one occurrence per sequence.

Raw read data are being processed for submission to ArrayExpress (accession numbers pending). The oligos used for the Nextera-based library prep are available in File S1, and the full list of CO intervals and midpoints is in File S2. File S3 is a list of genes in the 95th length percentile for regions without COs in our dataset. File S4 is an annotated list of NLR loci with a comparison of locus structure between Col and Ler.

## Results

### Development and validation of the Nextera LITE protocol

We developed a fast and inexpensive library preparation protocol, Nextera Low Input, Transposase Enabled (Nextera LITE), derived from the Illumina Nextera 24-sample reagent kit. Briefly, this protocol relies on dilution of the kit components, small reaction volumes, an optimal DNA template:tagmentation (TD) enzyme ratio, and optimized PCR amplification. For *A. thaliana*, we obtained fragment sizes suitable for the Illumina platform using 0.5 to 1 ng of DNA per 0.08 μL of TD enzyme (see Methods). Tagmented DNA fragments were directly amplified and dual-indexed in a PCR reaction without an intervening clean-up step, requiring less than two hours from start to finish and a reagent cost of $2.00 to $2.50 per library (Figure S1). We used a liquid-handling robot for several steps, but the entire protocol can also be easily performed by hand.

### Generating a high-density genome-wide CO map

The average read depth per library gave 1x to 2x genome-wide coverage per individual (Figure S2). Of the 1,920 Col x Ler F_2_ sequenced individuals, 95% had sufficient data quality and coverage to call COs. We added data from 363 previously analyzed Col x Ler F_2_ individuals (Choi *et al*. 2016; Underwood *et al*. 2018) that were produced using another library preparation method (Rowan *et al*. 2015, 2017). After processing and error correction for the entire data set (see Methods), we identified 17,077 COs in 2,182 individuals (File S2). We observed a mean of 7.8 COs per individual, which was consistent with previous estimates from Col x Ler and other crosses (Higgins *et al*. 2011; Salomé *et al*. 2012; Wijnker *et al*. 2013; Rowan *et al*. 2015). The median resolution for CO intervals – the distance between the closest scorable markers – was 1,102 bp, and three quarters of all CO positions could be estimated within an interval of 3 kb or less (Figure S3). We used the midpoints between flanking markers as estimates for CO positions in downstream analyses. In our dataset, the genetic distances in specific regions that had been previously been inferred by scoring recombination between fluorescent reporter genes in Col x Ler hybrids (Ziolkowski *et al*. 2015) were within 30% of the previously reported cM/Mb values, with the exception of a subtelomeric region on chromosome 2, which was 50% lower (Table S1). Since the recombination rate in this subtelomeric region is much higher in male than in female meiosis (Giraut *et al*. 2011), it is likely that this difference reflects measurement of both male and female meioses in our F_2_ population, while the distances estimated with fluorescent pollen markers reflect CO rates in male meiosis only (Ziolkowski *et al*. 2015). The distribution of CO rates in 200-kb windows along each of the five chromosomes (Figure 1A) was similar to what has been previously published for the species in general (Salomé *et al*. 2012) and for Col x Ler crosses specifically (Yelina *et al*. 2015; Choi *et al*. 2016), averaging 3.2 cM/Mb, with the highest rates in the pericentromeric regions and lowest rates in the centromeres.

**Figure 1.**
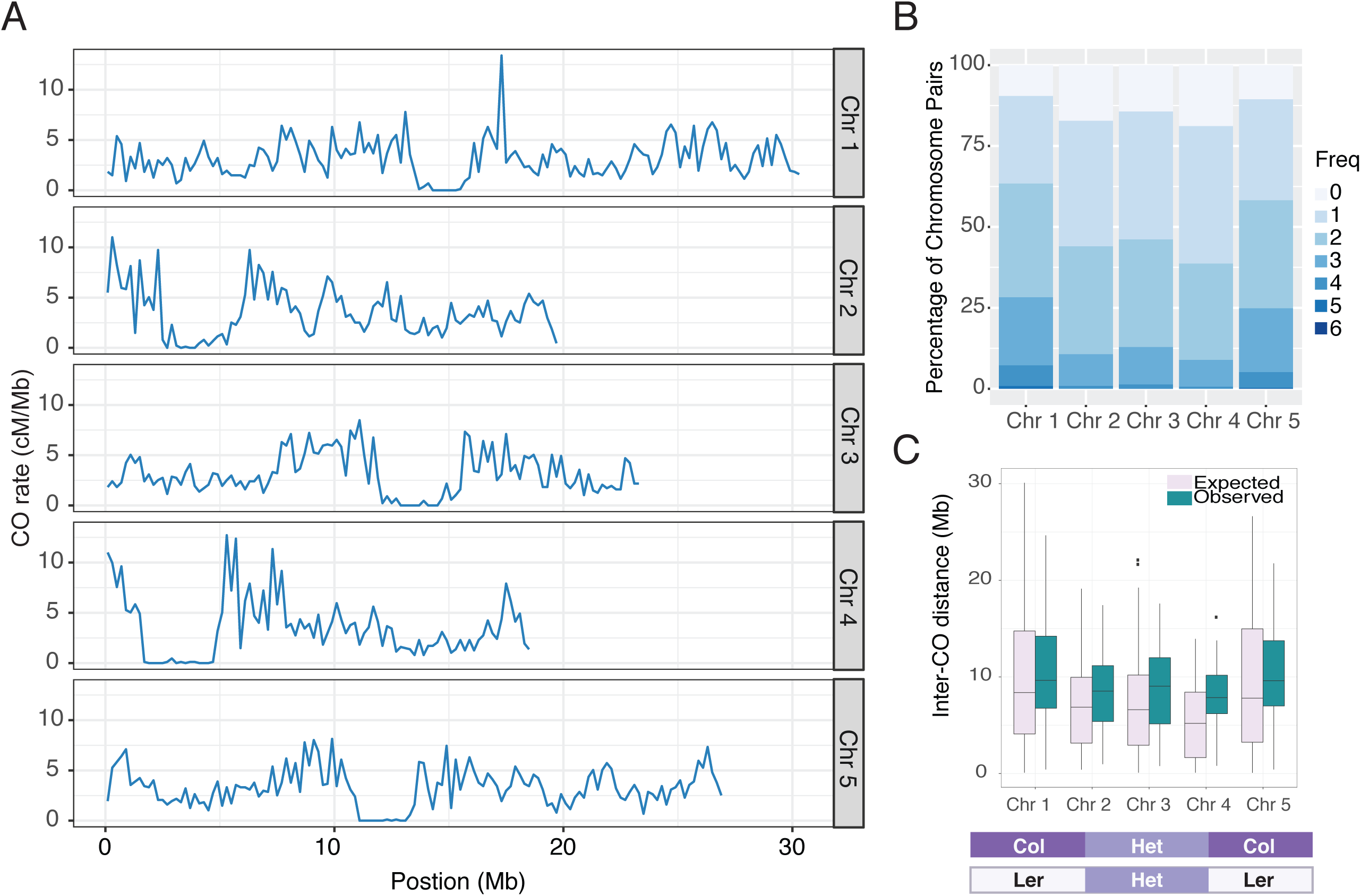
CO characteristics for a population of 2,182 Col X Ler F_2_ individuals. A) CO rates along all five chromosomes in fixed 200 kb windows. B) Frequency of CO numbers per chromosome pair for each chromosome. C) The distribution of inter-CO distances for chromosomes with cis double COs (determined by the genotype transitions shown in the schematic diagram below the plot). Observed distances are compared to the distances expected using randomly selected double CO sites.

The frequencies of COs per chromosome for all five chromosomes (Figure 1B) were similar to what has been reported previously (Salomé *et al*. 2012). The mean number of COs per chromosome pair ranged from 1.3 to 1.9 (Table 1) and was positively correlated with chromosome length (Figure S4, R^2^ = 0.96), agreeing with previous reports (Giraut *et al*. 2011; Salomé *et al*. 2012). To analyze COs that must have occurred on the same chromosome, we identified chromosome pairs with 2 or 3 COs (Figure 1C). Such double COs could only be unambiguously inferred when the sequence of blocks was Col-Het-Col or Ler-Het-Ler, and they therefore represent only a subset of all double COs. Using this approach, we found a total of 686 double COs with a mean inter-CO distance of 9,709,485 bp (Figure 1C, Figure S5), which was significantly greater than the expected mean of 8,770,824 bp for a matched set of random double COs (Figure 1C, p = 7.2 x 10^−7^, Mann-Whitney U-test), consistent with CO interference. The strength of interference was not different on different chromosomes (ANOVA, p = 0.47). The frequency of double COs was highest for the longest chromosomes, 1 and 5, which together accounted for 70% of all double COs that we could score (Table 1).

**Table 1.** COs per chromosome pair.

| Chr | Size (bp) | Total COs | Mean COs per pair | Double COs |
| --- | --- | --- | --- | --- |
| 1 | 30,427,671 | 4,153 | 1.9 | 260 |
| 2 | 19,698,289 | 3,021 | 1.4 | 78 |
| 3 | 23,459,830 | 3,191 | 1.5 | 76 |
| 4 | 18,585,056 | 2,825 | 1.3 | 49 |
| 5 | 26,975,502 | 3,887 | 1.8 | 223 |
| **TOTAL** |  | **17,077** |  | **686** |

### The effect of inversions on CO positions

Our dense CO dataset allowed us to examine in detail how SVs of different sizes affect local CO positioning. We first focused on the large paracentric inversion on chromosome 4 (Fransz *et al*. 2000, 2016; Zapata *et al*. 2016), relying on the flanking markers to establish whether a CO had occurred inside, since markers inside of SVs were filtered out during TIGER analysis. We did not find any COs within this ~1.2 Mb inversion (Figure 2A), compared with 162 expected COs in this region when a random distribution of COs was simulated (Table 2). The closest upstream CO occurred 5,425 bp from the left border and the closest downstream CO was just 1,245 bp from the right border, even though it was located at the edge of the centromere. The CO rate in the 200-kb region upstream of the border was considerably higher (4.6 cM/Mb) than the genome-wide average rate of 3.0 cM/Mb and lower (0.57 cM/Mb) in the 200-kb region downstream. The low CO rate in the downstream region is likely explained by its proximity to the centromere.

**Figure 2.**
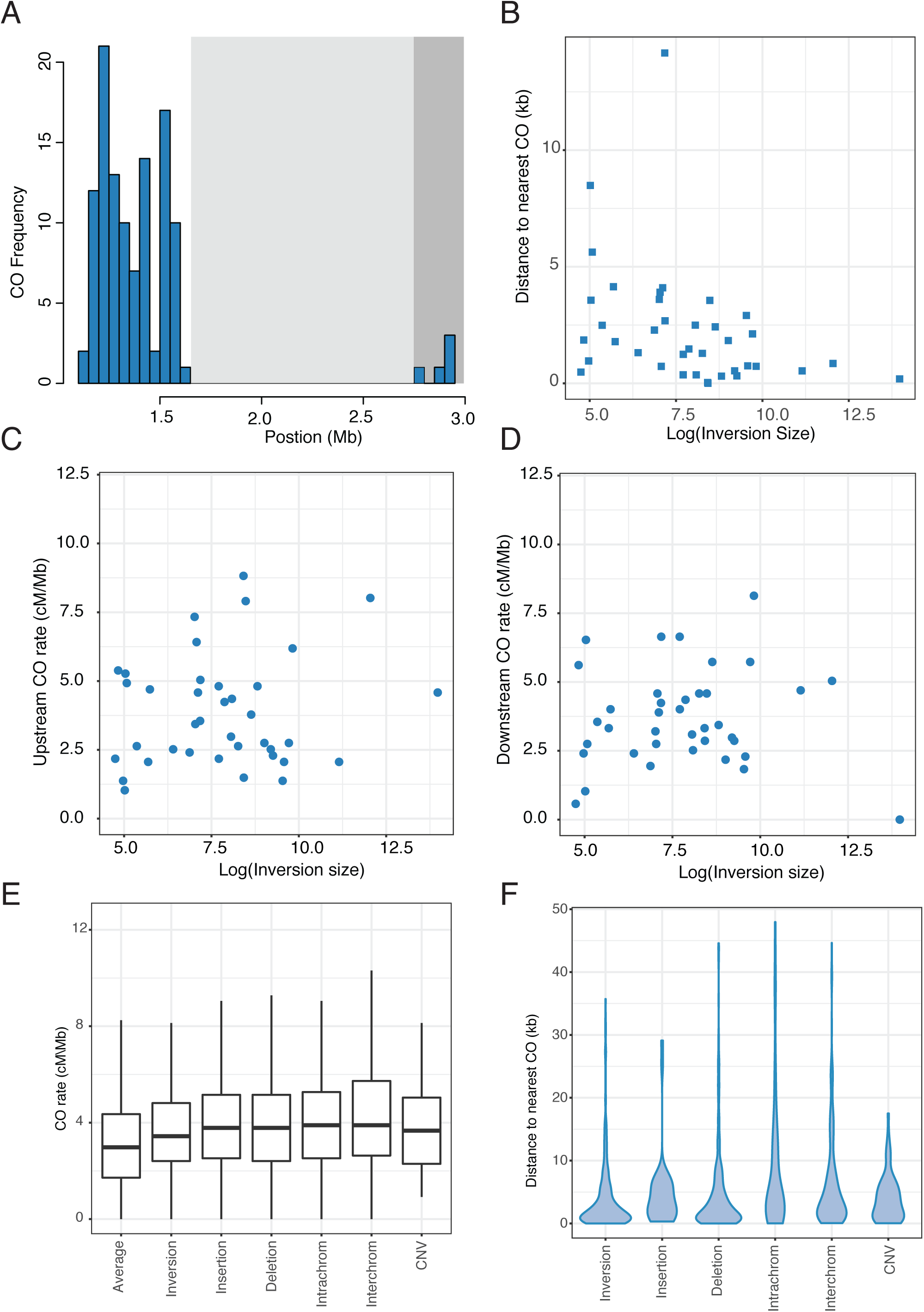
The effect of SVs on CO rates and positions. A) CO frequency in 50-kb bins in a region of chromosome 4 containing “the knob”, a 1.2-Mb inversion (light grey shaded region) that is adjacent to the centromere (dark grey shaded region). B) Minimum CO distance from the borders of all inversions in relation to the log10 of the inversion size. C) CO rates (cM/Mb) in the 200 kb upstream of inversion borders. D) CO rates (cM/Mb) in the 200 kb downstream of inversion borders. E) CO rates (cM/Mb) in the 200 kb up- and downstream of the borders for inversions, insertions, deletions, transpositions (intrachrom) and translocations (interchrom) and copy number variations (CNV). F) Distances to the nearest CO for inversions, insertions, deletions, transpositions (intrachrom) and translocations (interchrom) and copy number variations.

**Table 2:**
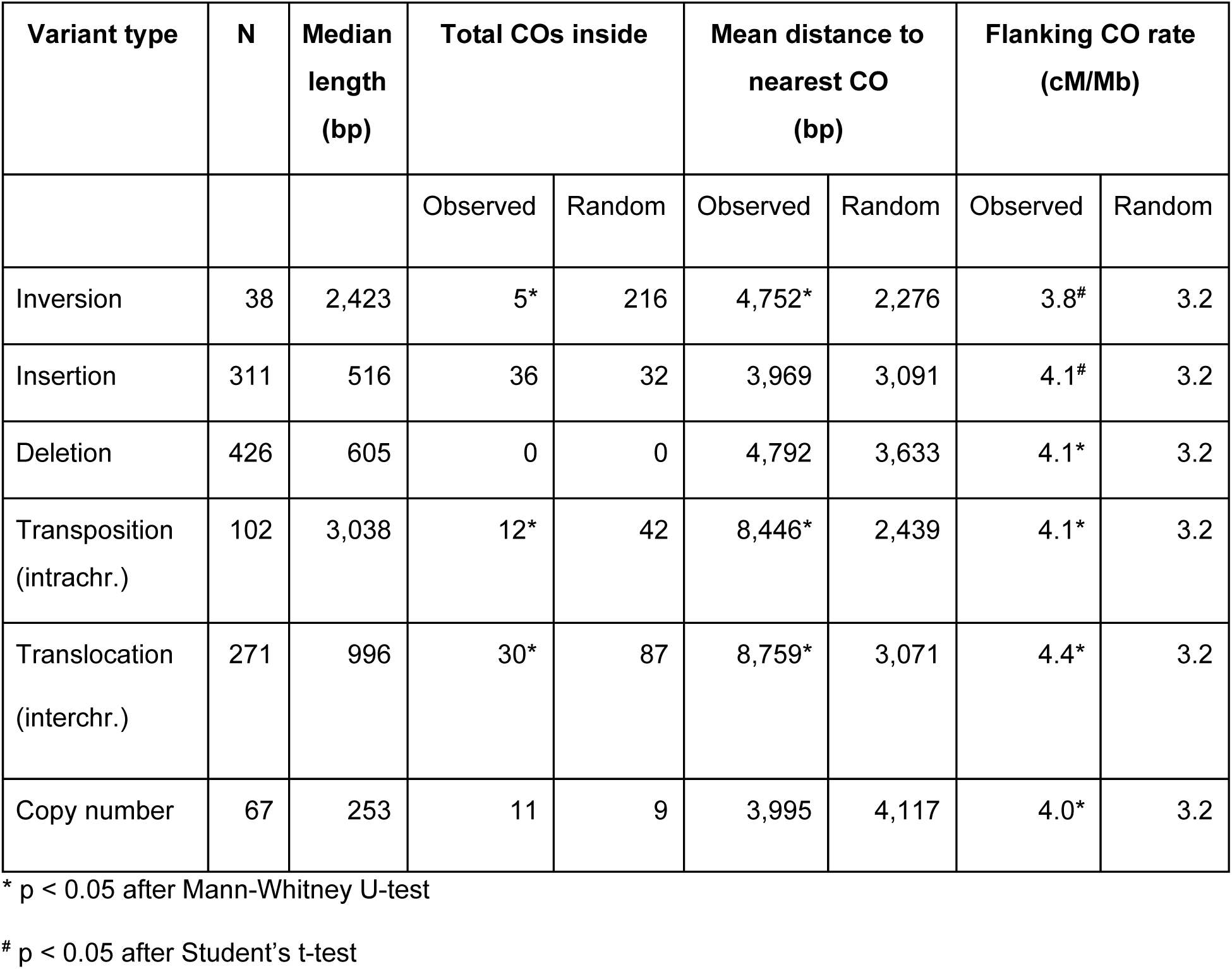
COs within and next to SVs.

| Variant type | N | Median length (bp) | Total COs inside |  | Mean distance to nearest CO (bp) |  | Flanking CO rate (cM/Mb) |  |
| --- | --- | --- | --- | --- | --- | --- | --- | --- |
|  |  |  | Observed | Random | Observed | Random | Observed | Random |
| Inversion | 38 | 2,423 | 5* | 216 | 4,752* | 2,276 | 3.8<sup>#</sup> | 3.2 |
| Insertion | 311 | 516 | 36 | 32 | 3,969 | 3,091 | 4.1<sup>#</sup> | 3.2 |
| Deletion | 426 | 605 | 0 | 0 | 4,792 | 3,633 | 4.1* | 3.2 |
| Transposition (intrachr.) | 102 | 3,038 | 12* | 42 | 8,446* | 2,439 | 4.1* | 3.2 |
| Translocation (interchr.) | 271 | 996 | 30* | 87 | 8,759* | 3,071 | 4.4* | 3.2 |
| Copy number | 67 | 253 | 11 | 9 | 3,995 | 4,117 | 4.0* | 3.2 |

\* p < 0.05 after Mann-Whitney U-test
<sup>#</sup> p < 0.05 after Student's t-test

We also did not find any COs inside another large paracentric inversion on chromosome 3 (170 kb). Flanking COs were relatively nearby, within 10 kb on either side of the inversion breakpoints (Figure S6). The 200 kb upstream and downstream regions of this 170 kb inversion had CO frequencies of 8.0 and 5.0 cM/Mb.

To examine CO patterns more broadly, we looked at the overlap between CO intervals and 47 inversions of different sizes (range 115 - 1,178,806 bp; median: 2,221 bp; (Zapata *et al*. 2016)). Because COs inside paracentric inversions lead to the formation of dicentric and acentric products (McClintock 1931, 1933), which may be lost or lead to aneuploidy, COs inside paracentric inversions are expected to produce fewer viable offspring. A total of 13 inversions overlapped with CO intervals, which was significantly lower than expected by chance (p=0.0002, established with 5,000 permutations). To further assess the impact of inversions on CO positions at a local scale, we excluded the centromeric regions where effects of inversions would be masked by the inherently low recombination rate. In a final set of 38 non-centromeric inversions and 16,776 genome-wide COs, we found only five events where the mid-point CO estimate was within an inversion, which was significantly different from 216 internal COs when a random distribution was simulated (p = 0.004, Mann-Whitney U test). Moreover, the CO intervals spanning these five inversions were 13 kb on average, meaning that the resolution of these CO positions was much lower than the median CO resolution of the dataset. Thus, it is likely that these COs occurred outside of the inversion boundaries. There was no effect (R^2^ = 0.0006, p = 0.89) of inversion size on the distance to the nearest CO (Figure 2B) and the mean distance of an inversion breakpoint to the closest CO midpoint was 4,752 ± 957 bp (mean ± sem), which was significantly greater than the mean distance when a random CO distribution was simulated (2,276 ± 430 bp, p = 0.007, Mann-Whitney U test). The mean CO rate in the 200 kb up- and downstream of the inversions (3.8 ± 0.26) was significantly higher (p = 0.03, Student’s t test) than the rates in a random comparison (3.2 ± 0.08). The observed CO rates in the flanking 200-kb regions of inversions (Figure 2C,D) did not depend on inversion size (R^2^ = 0.01346, p = 0.49). We conclude that COs are suppressed within inversions of all sizes, but this suppression does not extend far beyond the inversion borders (< 10 kb).

### The effect of other SVs on CO positions

To determine whether the observed effects of inversions on COs are also seen with other SV types, we compared locations of COs in relation to insertions, deletions, tandem copy number variants (CNVs), transpositions, and translocations (Zapata *et al*. 2016). For all SV types except tandem CNVs, the overlap between CO intervals and the regions containing SVs was significantly lower than expected by chance (Table S2). When looking at CO positions, only transpositions and translocations had fewer internal COs and an increased distance to the nearest CO than expected from a matched random distribution of COs (Table 2). The distance to the nearest crossover for all SV types was not affected by the SV length (Figure S7). CO rates in the flanking 50, 100, or 200 kb up- and downstream of all SVs were higher than both the observed genome-wide average and a matched random distribution of COs (Figure 2E, Table 2, Figure S8, and Table S3) and independent of the size of the variant region (Figure S9). We conclude that there is a general trend of local suppression of COs within SVs of different types and lengths, which occasionally extends several kb beyond the SV borders.

### COs, double-strand breaks, and SVs

Meiotic COs occur through repair of programmed double strand breaks (DSBs) generated by SPO11 complexes during prophase I of meiosis (Szostak *et al*. 1983; Keeney *et al*. 1997; de Massy 2013). We therefore sought to examine whether suppression of COs around SVs was associated with reduced DSB formation in those regions. Meiotic DSBs have been mapped in *A. thaliana* via purification and sequencing of SPO11-1-oligonucleotides, which mark DSB sites (Choi *et al*. 2018). These data were used to define 5,914 SPO11-1-oligo hotspots (Choi *et al*. 2018), which we compared with CO positions and the locations of SVs throughout the genome (Figure 3A). As expected, the overlap between SPO11-1-oligo hotspots and CO intervals was significantly greater than random (Figure 3B, p = 0.0002 after 5000 permutations). Importantly, the overlap between SPO11-1-oligos and SVs was not significantly different from that expected by chance, indicating that the initiation of COs is not different inside and outside SVs (Figure 3C, p=0.09 after 5,000 permutations). The density of SPO11-1-oligo hotspots and COs were also positively correlated in 200-kb windows across the genome (Figure 3D, R^2^ = .34, p < 2 x 10^−16^). Over half of all COs (53%) were within 5 kb of a SPO11-1-oligo hotspots (Figure 3E). Previous work established a relationship between low nucleosome occupancy and elevated levels of SPO11-1-oligonucleotides (Choi *et al*. 2018). Consistent with this, we observed that CO midpoints were associated with reduced nucleosome occupancy, measured via micrococcal nuclease sequencing (Choi *et al*. 2016), and elevated SPO11-1-oligonucleotides (Figure 3F,G). Taken together, these results suggest that suppression of COs around SVs occurs downstream of DSB formation.

**Figure 3.**
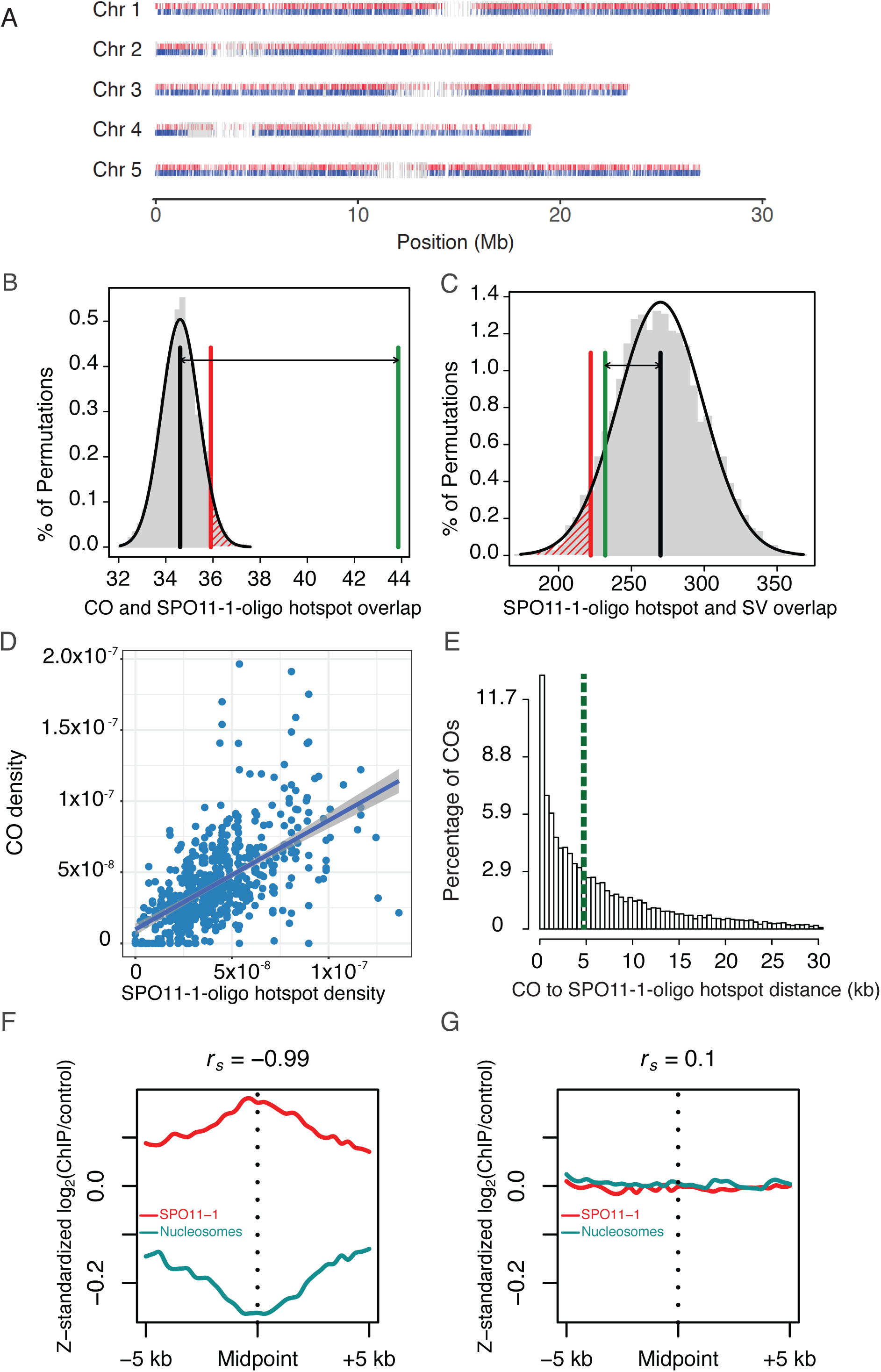
Coincidence of COs, SPO11-1-oligo hotspots and SVs. A) Locations of SPO11-1-oligo hotspots, COs, and SVs or each chromosome. The position of the midpoint of the SPO11-1 oligo hotspot is shown in red, the CO midpoints in blue and the locations of SVs in gray. Permutation tests were performed to test for overlaps between (B) SPO11-1-oligo hotspots and CO intervals and (C) SPO11-1-oligo hotspots and SVs. Vertical black lines indicate the mean number of overlaps expected in 5000 random permutations. The vertical red lines indicate the number of overlaps where p = 0.05. The vertical green line indicates the observed number of overlaps. The double arrow highlights the difference between the mean of 5,000 permutations and the observed number. D) Correlation between the percentage of SPO11-1-oligo hotspots and the percentage of COs in 200-kb windows. The equation of the line is 0.77*x* + 0.002, R^2^ = 0.34, p < 2 x 10^−16^. E) Histogram of distances from CO breakpoints to the border of the nearest SPO11-1 oligo hotspot. The vertical green dashed line indicates the median. Outliers representing the furthest 5% of distances are not shown in the plot. F) SPO11-1-oligo (red) or nucleosome occupancy (blue, MNase-seq) (z-score standardized log_2_(signal/input)) in 10 kb windows around CO midpoints. G) As for F), but showing analysis of the same number of randomly chosen positions.

### Regions devoid of COs (“CO deserts”)

With our ultra-dense CO data, we were able to identify large regions of the genome where CO occurrence was much lower than expected by chance. The median distance between all COs was 2,610 bp and over 80% of the COs were spaced within 10 kb of another CO in the dataset. We selected the largest 5% of inter-CO distances (minimum 25,563 bp and maximum 1,419,083 bp, n=839) and investigated these “CO deserts” to uncover genomic features that suppress CO formation. We expected that these regions would correspond to SVs, but the overlap between CO deserts and all SVs was not significantly different from that expected by chance (p=0.1 after 5,000 permutations, Figure S10A). While only 6.3% of CO deserts were located within centromeres, these were among the longest CO deserts (top 2%). CO deserts were significantly enriched for DNA methylation at CG sites even when centromeres were excluded (Figure S10B, p < 2.2e-16 with Student’s T-test or Mann-Whitney U-Test). Although SVs were not more likely to be found in CO deserts, they were significantly enriched for CG DNA methylation (Figure S10C, p < 2.2e-16 with Student’s T-test or Mann-Whitney U-Test). Over 50% of the genes in CO deserts consisted of transposable elements (Supplemental File 3). Other genes that were enriched in CO deserts were those involved in meiosis, mitosis/cell-cycle, and DNA repair/metabolism (Table S4). We conclude that CG DNA methylation is associated with CO suppression in SVs, TEs, and some regions of the genome that contain protein coding genes.

### COs and NLR gene clusters

Since COs reshuffle existing genetic diversity, they represent an important mechanism for generating alleles with new functions. This is especially important for nucleotide binding site leucine rich repeat (NLR) genes involved in disease resistance, as COs have been implicated in the generation of new resistance specificities in plants (Richter *et al*. 1995; Parniske *et al*. 1997; McDowell *et al*. 1998; Noël *et al*. 1999). Many NLR genes in the *A. thaliana* genome are found in clusters that exhibit structural variation, which may locally suppress CO formation (Chin *et al*. 2001; Meyers *et al*. 2003; Alcázar *et al*. 2014; Chae *et al*. 2014; Van de Weyer *et al*. 2019). Indeed, a previous study showed that NLR genes in Col x Ler hybrids had variable CO rates with some genes overlapping CO hotspots while others, including those with a high degree of structural variation, were CO coldspots (Choi *et al*. 2016). The dense CO dataset generated in the current study enabled a deeper examination of the patterns of COs within and around NLR genes and other genes involved in defense.

The number of overlaps between CO intervals and NLR/other defense genes was not significantly different from that expected by chance (p=0.23, 5,000 permutations, Figure S11). Considering CO positions, we found that 55 of 197 of defense-related genes (28%) contained one or more COs, which was not significantly different from randomized positions (p = 0.06, Wilcoxon test). However, the mean distance from defense gene locus borders to the nearest CO was 7,107 bp, which was significantly further away than expected from a random CO distribution (2,212 bp, p = 7.5 X 10^−6^, Wilcoxon test). CO rates in the flanking 200 kb up- and downstream were slightly, but not significantly elevated.

We categorized NLRs by their locus structure (Figure 4A), and found CO hotspots and coldspots among genes of all locus types. Singleton NLR genes and those in loci with inverted and tandem repeat structures trended toward higher proportions of CO coincidence compared with complex and tandem inverted locus structures (Figure 4B, Table 3). When Col and Ler have the same NLR locus structure (Figure 4C, File S4), the CO rates are generally higher than in NLR loci where the two parents differ in their structure (Figure 4D). However, the number of overlaps between NLR genes and CO intervals is not significantly different between the two classes (Figure S11B,C).

**Figure 4.**
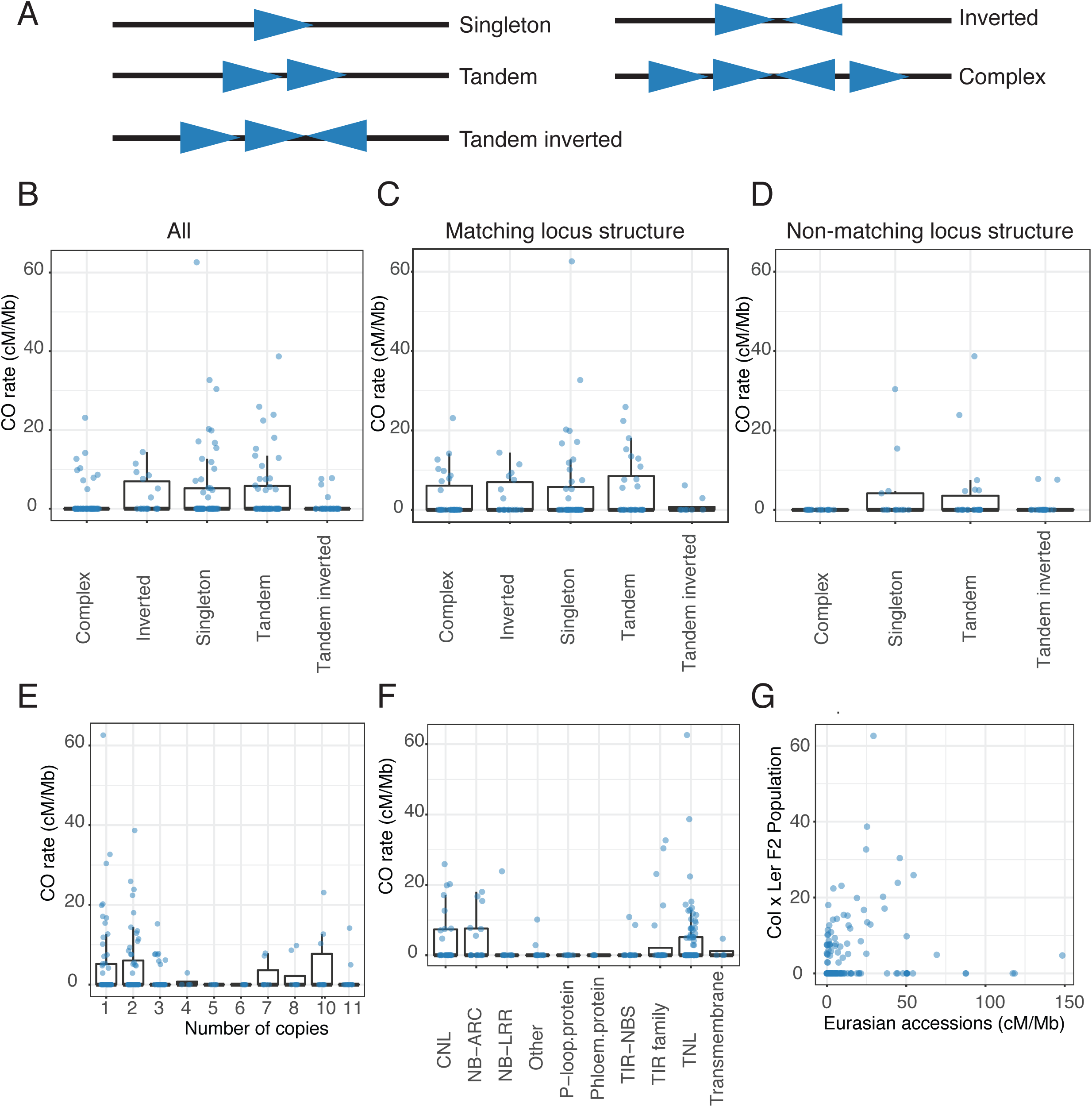
CO frequency in disease resistance genes. A) Schematic drawings representing different locus structures for nucleotide binding site leucine rich repeat (NLR) and other disease resistance genes. B) CO rates for all NLR loci and other disease resistance genes (File S4) based on Col locus structure. C) CO rates for NLR loci with similar structure in Col and Ler. D) CO rates for NLR loci that differ in structure between Col and Ler. E) CO rates for all NLR loci as a function of copy number at the locus for Col (see Figure S12 for Col/Ler differences). F) CO rates for NLR loci based on Col annotation. G) CO rates in NLR genes in our Col x Ler F2 population compared with the historical recombination rate across Eurasian accessions.

**Table 3:** COs within NLR genes as a function of different structural classes.

| Locus Type | Genes | Number with COs | Proportion with COs |
| --- | --- | --- | --- |
| Complex | 47 | 9 | 0.19 |
| Inverted | 18 | 7 | 0.39 |
| Singleton | 57 | 18 | 0.32 |
| Tandem | 50 | 18 | 0.36 |
| Tandem inverted | 25 | 4 | 0.16 |

COs trended toward being more prevalent in loci with only one or two genes (Figure 4E), especially when both Col and Ler had the same number of copies in a locus (Figure S12). COs showed a general trend toward exhibiting higher levels in TIR domain NLRs (TNLs), coiled-coil domain NLRs (CNL), and those with only nucleotide-binding (NB-ARC) domains than in other categories of NLR genes (Figure 4F, Table S5). There was no correlation between historical recombination rates among Eurasian accessions (Choi *et al*. 2016) and CO rates within NLR genes among Col x Ler F_2_ individuals (Figure 4G). Locations of CO positions in a focal set of NLR genes that were shown to have high CO rates (Figure S13) were generally consistent with a previous study (Choi *et al*. 2016).

### COs, chromatin and sequence features

COs tend to be associated with several hallmarks of open chromatin in plants, including low levels of DNA methylation, high levels of histone H3K4me3 methylation, and low nucleosome density (Liu *et al*. 2009; Yelina *et al*. 2012; Marand *et al*. 2017; Choi *et al*. 2018). COs have been found to be enriched close to the 5′ and 3′ flanking regions of genes, at transcription start sites, *cis*-regulatory regions, and regions where the histone variant H2A.Z has been deposited (Choi *et al*. 2013; Wijnker *et al*. 2013; Marand *et al*. 2017), which is consistent with our finding that COs were likely to be in nucleosome-depleted and SPO11-1-oligo enriched regions of the genome (Figure 3 G,H). Therefore, we expected to observe that COs would overlap with DNase I hypersensitive (DNase I HS) sites and Assay for Transposase Accessible Chromatin sequencing (ATAC-seq) sites, which are two independent indicators of regions of open chromatin/cis-regulatory elements (Sullivan *et al*. 2014; Maher *et al*. 2018). We found that our CO intervals showed a significant overlap with DNaseI HS sites (Figure 5A). 4,338 of 17,077 CO midpoints (~25%) occurred in a DNase I HS site and a further 36% of COs occurred within 1 kb up- or downstream of a DNase I HS site. When a random distribution of COs was simulated, only 8% occurred within a DNase I HS site and 20% within 1 kb of a DNAse I HS site. Over 95% of COs occurred within 6 kb of a DNase I HS site (Figure 5B), with a median distance of 558 bp to the nearest DNase I HS site. In comparison, 95% of a random set of COs were within 42 kb of a DNase I HS site and the median distance was 1 kb. Results were similar for ATAC-seq sites, where 95% of COs were within 6.5 kb of an ATAC-seq site (Figure S14), compared with 95% of a matched set of random COs occurring within 39 kb of an ATAC-seq site.

**Figure 5.**
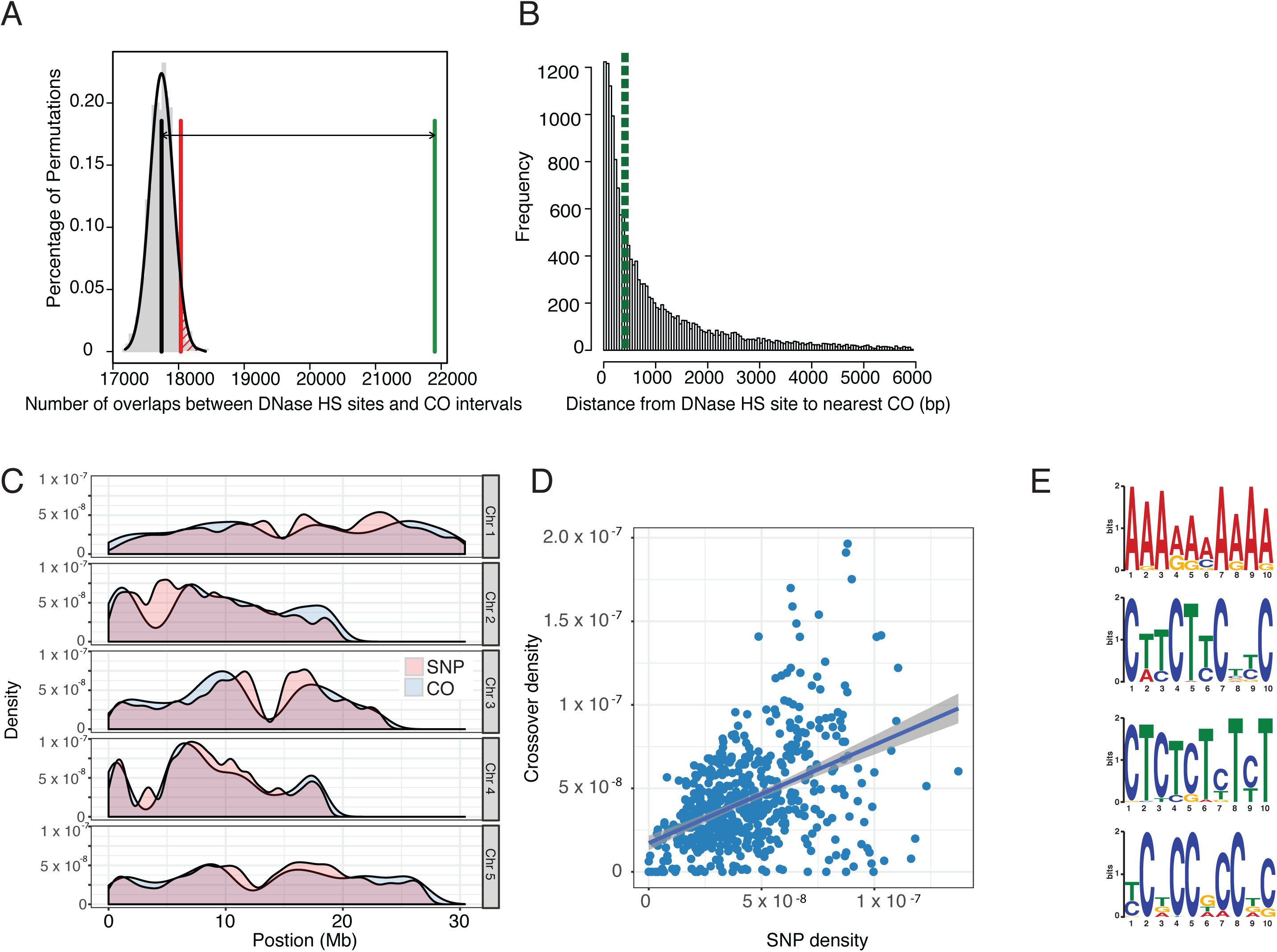
Chromatin and sequence features associated with COs. A) Permutation tests of overlaps between DNase hypersensitive (HS) sites and CO intervals. Vertical black line indicates mean number of overlaps expected in 1,000 random permutations. Vertical red line indicates the number of overlaps where p = 0.05. Vertical green line indicates the observed number of overlaps. Double arrow highlights the difference between the expected mean and the observed number. B) Distribution of distances from CO breakpoints to the nearest DNase HS site border. The vertical dashed green line indicates the median. The top 5% of distances are omitted from this plot for ease of visualization. C) Overlay of CO and single nucleotide polymorphism (SNP) density along all five chromosomes. D) Correlation between CO density and SNP density in 200-kb fixed windows. Line indicates best fit and grey shaded area represents the 95% confidence interval. The equation of the line is 5.872e-01*x + 1.726e-08. R^2^ = 0.21. p < 2.2e-16. E) Sequence motifs enriched in the 500 bp up- and downstream of the CO breakpoint site for a subset of crossovers (1,000 per chromosome). The top four detected motifs are shown.

Our dataset also allowed us to reexamine the question of whether recombination is itself mutagenic and would therefore be associated with genomic polymorphism (Cao *et al*. 2011; Long *et al*. 2013; Ziolkowski *et al*. 2015; Yang *et al*. 2015). However, we also note that divergence between homologs suppresses recombination (George and Alani 2012; Liu *et al*. 2012; Chakraborty and Alani 2016), as we observed within SVs (Figure 2). Thus, one might expect that these two opposing forces would lead to differences in the correlation between SNPs and COs over evolutionary time. We found that the pericentromeres are regions of relatively high SNP and CO density for the Col/Ler cross, despite also containing higher levels of heterochromatin (Figure 5C Figure S15A, (Yelina *et al*. 2012; Choi *et al*. 2013, 2018; Underwood *et al*. 2018)). The densities of Col/Ler SNPs and COs among all 200-kb windows across the genome showed a moderate, but statistically significant positive correlation (R^2^ = 0.21, Figure 5D), similar to that reported previously (Serra *et al*. 2018; Fernandes *et al*. 2018). In addition, the pericentromeres have been previously shown to have a general AT sequence bias (Wijnker *et al*. 2013). At the fine scale, several DNA sequence motifs have been associated with COs in Col x Ler crosses, including a polyA/T motif, a CTT/GAA repeat (Choi *et al*. 2013; Wijnker *et al*. 2013; Shilo *et al*. 2015) and a CCN repeat (Shilo *et al*. 2015). Therefore, we used MEME to identify motifs in the 500 bp up- and downstream of COs in random subsets of 500 and 1,000 COs per chromosome. We found all three of these motifs (Figure 5E) in our CO data, plus a fourth motif (CT repeat). The polyA and CTT/GAA repeat motifs were the most common - nearly 90% of COs were within 500 bp of one or the other of these repeats (Table 4 and Table S6), while the CCN and CT repeat motifs were found in a minor fraction of the analyzed COs. Taken together, these data indicate that COs between Col and Ler are strongly associated with these four sequence motifs.

**Table 4:** Sequence motifs in 1-kb regions spanning of a random subset of 5,000 COs.

| Motif | Occurrence | Percent of all COs |
| --- | --- | --- |
| PolyA/T | 4,314 | 86.2 |
| CTT/GAA | 1,527 | 30.5 |
| CT/GA | 1,721 | 34.4 |
| CCN/GGN | 901 | 18.0 |
| PolyA/T or CTT/GAA | 4,492 | 89.8 |
| All four motifs combined | 4,643 | 93.8 |

## Discussion

We employed a new low-cost rapid genomic DNA library prep to sequence the genomes of over 1,800 F_2_ hybrid individuals from a cross between two commonly used *A. thaliana* accessions, and to map COs at a very high density and precision. The availability of high-quality reference genomes for each of these accessions (Arabidopsis Genome Initiative 2000; Lamesch *et al*. 2012; Zapata *et al*. 2016) enabled a detailed analysis of the effects of SVs on CO formation and a fine-scale examination of COs in relation to other sequence and chromatin features.

### An improved method for genomic DNA library construction

Our library preparation method had a success rate of around 95% when processing hundreds of samples in under 3 hours. Furthermore, at a cost of ~$2.50 per sample, this method is, as of writing this manuscript, highly cost effective and compares favorably in terms of speed with similar protocols (Baym *et al*. 2015; Pisupati *et al*. 2017). A version of this protocol could also be used with homemade Tn5 transposome and buffer (Hennig *et al*. 2018). Combining the COs determined from 1,829 F_2_ hybrid genomes with previously published CO data for the same cross (Choi *et al*. 2016; Underwood *et al*. 2018), we obtained a final dataset of 17,077 COs. The pattern of CO rates along the chromosomes (Figure 1) was in good agreement with previously reported genome-wide CO data generated with a variety of approaches (Giraut *et al*. 2011; Salomé *et al*. 2012; Yelina *et al*. 2012; Wijnker *et al*. 2013). From extracted DNA to final CO data, a genetic map could be built in about 1.5 weeks for hundreds of samples using this combination of library preparation and CO calling with TIGER analysis (Rowan *et al*. 2015).

### Insights into the mechanism for CO suppression inside SVs

Given the high density of COs in our dataset, we were able to examine the relationship between COs and multiple types of SVs in fine detail. Observed COs were suppressed inside inversions and transpositions, and were slightly elevated in the flanking regions of all SV types (Figure 2, Table 2, Table S2, Table S3, Figure S8). One interpretation for the elevated flanking CO rates is that the lack of COs inside the SVs is compensated by additional COs in the flanking regions. However, such a simple scenario would predict that larger SVs would have higher flanking CO rates, but this is not what we found (Figure S9). A more likely explanation is that SVs occur more often in regions that are already prone to high rates of recombination, as recombination is a mechanism for generating SVs (George and Alani 2012; Liu *et al*. 2012). In other words, COs in the vicinity of SVs are not increased because of SVs, but SVs are more likely to occur in regions with elevated CO rates. For inversions and translocations, the suppression of COs within the variant regions tends to extend moderately beyond the borders of the variant regions, as the distance to the nearest CO is slightly further than expected under a random CO distribution (Table 2), although this was not correlated with SV length (Figure S7). This is in contrast to *Drosophila* interspecific crosses, where suppression of recombination extends over 2 Mb beyond the boundaries of inversions of different sizes (Kulathinal *et al*. 2009; Stevison *et al*. 2011). Below, we discuss five possible mechanisms explaining why COs are less likely to occur inside of SVs: *(1)* DSB formation in variant regions is reduced; *(2)* reduced sequence similarity means that there is no substrate for recombinational repair; *(3)* COs that form within SVs create inviable gametes and thus cannot appear in progeny; *(4)* variant regions are physically prevented from interacting with the homologous chromosome by the structure and organization of the synaptonemal complex; and *(5)* DNA methylation suppresses COs in SVs.

1. Possible mechanism: DSB formation in variant regions is reduced. SPO11-1-oligo hotspots are as common in SVs as outside SVs (Figure 3), suggesting that meiotic DSBs that occur in these regions are repaired as non-COs if the homologous chromosome is used, or they are repaired using the sister chromatid. A caveat is that SPO11-1-oligos hotspots were mapped only in Col homozygotes (Choi *et al*. 2018), and there is a possibility that these hotspots are distributed differently in Ler and/or Col/Ler F_1_ hybrids. A recent study of inversion heterozygotes in *Drosophila melanogaster* crosses suggested that the DSB number is similar in such regions, but that they are preferentially repaired as NCOs (Crown *et al*. 2018). While we cannot rule out a role for suppression of COs by prevention of DSBs within SVs in *A. thaliana*, it is likely not a major contributing factor.
2. Possible mechanism: reduced sequence similarity means that there is no substrate for recombinational repair. SV types differ in their level of sequence divergence. In the case of inversions, sequence identity between the two chromosomes means that pairing is possible, but the portion of the chromosome containing the inversion must be looped so that the region is in the same orientation on both homologues. Insertion/deletion (indel) polymorphisms and transpositions have no direct counterpart on the other chromosome, but the indel sequence could contain repetitive elements that occur in nearby non-allelic loci and provide a substrate for homologous recombination. Tandem CNVs should provide sufficient homology for recombination. Because inversions and tandem CNVs have homology, but exhibit reduced CO rates, it suggests that a lack of homology alone is not a major factor suppressing COs in these regions.
3. Possible mechanism: COs that form within SVs create inviable gametes and thus cannot appear in progeny. Although inversion loops would facilitate the alignment of homologous regions, a crossover that occurs through this process is expected to generate a dicentric bridge and an acentric chromosome. This could potentially lead to inviable gametes due to aneuploidy and explain why COs in inversions are not observed in F_2_ progeny. However, a previous cytological examination of male meiotic prophase I in Col x Ler hybrids did not reveal inversion loops, and dicentric bridges were rarely observed in anaphase I (Ji 2014). In hybrids of wild and domesticated maize hybrids that were heterozygous for a ~50-Mb inversion, only 7 of 167 male meiocytes showed dicentric bridges, and pollen viability and seed set were normal (Fang *et al*. 2012). This suggests that there are mechanisms to prevent COs that would generate defective gametes (at least in the case of inversions), but the frequency of deleterious COs in SVs in gametes needs to be assessed before a conclusion can be reached.
4. Possible mechanism: variant regions are physically prevented from interacting with the homologous chromosome by the structure and organization of the synaptonemal complex. The formation of COs occurs within a highly-organized protein-DNA structure known as the synaptonemal complex (SC), consisting of a central element flanked by two lateral elements (Zickler and Kleckner 1999). This structure of this complex is highly conserved across a broad taxonomic range and functions to hold homologous chromosomes in proximity during CO formation. Recombination is thought to occur between the homologues in proximity to the SC central element, where proteins that facilitate the repair of DSBs in favor of a CO outcome, which is marked by MLH1 foci (Bogdanov *et al*. 2007). For each homolog, the rest of chromosome that is not participating in crossing over is organised as chromatin loops tethered to the central and lateral elements, known as tethered-loop/axis architecture (Blat *et al*. 2002). It is possible that the portions of the chromosomes that contain SVs are prevented from interacting with the central element and the homologous chromosome. This might explain why in some cases, the distance to the nearest CO was slightly further away than expected by chance. Future research aimed towards obtaining fine-scale organization of the DNA sequences bound to the SC will help clarify whether this is a mechanism contributing to suppression of COs in SVs.
5. Possible mechanism: DNA methylation prevents COs from occurring in SVs. The level of CG DNA methylation was higher in 10-kb windows that overlapped with SVs in our analysis than in 10-kb windows that did not overlap SVs (Figure S10C). We also found that regions surrounding SVs had elevated CO rates (Figure 2E). We propose that newly arising SVs are likely to occur via recombination; once generated, they have a probability of becoming methylated. If DNA methylation in SVs prevents COs that are otherwise deleterious from occurring, then this could select for methylation around SVs. The mechanism by which DNA methylation prevents crossing over could act through organization of the SC (as proposed in *(4)* above) or through other mechanisms, such as the promotion of non-crossover DNA repair pathways.

### Other DNA sequence features associated with COs

With this dense CO dataset, we defined an even more nuanced and detailed picture of CO distribution in relation to NLR and other defense genes (Choi *et al*. 2016), chromatin accessibility (Liu *et al*. 2009; Choi *et al*. 2013; Yelina *et al*. 2015; Shilo *et al*. 2015; Marand *et al*. 2017) and DNA sequence motifs (Choi *et al*. 2013; Wijnker *et al*. 2013; Shilo *et al*. 2015) at the fine scale. Recombination has an important role in generating sequence diversity in NLR genes over evolutionary time (Parniske *et al*. 1997; Parniske and Jones 1999; Noël *et al*. 1999; Sun *et al*. 2001; Wulff *et al*. 2004), and it can alter resistance specificites in experimental crosses either by generating chimeric genes (Collins *et al*. 1999), or by altering locus copy number (Chin *et al*. 2001). Thus, it might be expected that CO rates in NLR genes would be high. However, when examining NLR genes genome-wide in an experimental cross, NLR genes were found to be equally likely CO hotspots or coldspots, and this was in some cases related to variation in locus architecture (Choi *et al*. 2016). Roughly half of the NLR genes in the *A. thaliana* genome exist in clusters with one or more paralogs (Meyers *et al*. 2003). In this study, we found that tandem and inverted duplications and singleton NLR genes had the highest proportion of genes overlapping COs (Figure 4D and Table 3). The proportion of genes with COs was generally lower for loci that differed in structure between Col and Ler. Complex loci, defined as having at least four paralogs, only featured COs when Col and Ler had the same locus structure. CO proportions and rates were highest in loci where Col and Ler both had either only one or only two NLR gene copies. Recombination can therefore not only generate diversity in NLR genes that may alter resistance specificity, but it may also generate structural variation that suppresses recombination over time, potentially leading to resistance supergenes where several disease resistance specificities are inherited together via linkage. This would also protect newly emerged NLR genes that have not yet required a function from elimination, as long as there is a positively selected gene in such a cluster.

Methylation of histone H3 lysine 4 (H3K4me3), a hallmark of actively transcribed genes is positively associated with COs in plants (Yelina *et al*. 2012, 2015; Marand *et al*. 2017), while dense DNA methylation, an indicator of repressed chromatin state, is negatively associated with COs (Yelina *et al*. 2012, 2015; Rodgers-Melnick *et al*. 2015; Marand *et al*. 2017). Chromatin accessibility, as determined by DNase I hypersensitivity, is positively associated with COs in potato (Marand *et al*. 2017) and maize (Rodgers-Melnick *et al*. 2015). Consistently, we also found COs to be highly associated with DNase I hypersensitivity (Figure 5A,B) and with another measure of chromatin accessibility, ATAC-seq sites (Figure S14). Three previously described CO-associated sequence motifs, poly-A, CTT and CCN repeats (Wijnker *et al*. 2013; Yelina *et al*. 2015; Melamed-Bessudo *et al*. 2016) are found in regions of open chromatin (Shilo *et al*. 2015). At least one of these motifs, in addition to a CT repeat, was present within 1 kb of nearly all COs that were examined (Figure 5E, Table 4).

## Conclusions

The ultra-dense catalog of COs in this study has extended and refined our knowledge of various factors that influence the position and frequency of COs along *A. thaliana* chromosomes. We found that all types and sizes of SVs can affect local CO positioning, not only the large-scale rearrangements of the type that have been more extensively studied in the past. Since the suppressive effect of smaller SVs does not spread far beyond the borders, only SVs that are large enough to contain multiple genes are expected to have the potential to create supergenes containing blocks of jointly inherited favorable alleles.

## Author contributions

B.R., D.W. and I.H. designed the study. D.H. developed the Nextera LITE library preparation protocol. B.R. and T.F. grew all of the plants and performed all of the molecular biology work. B.R., A.T. and I.H. analyzed the data and B.R. wrote the manuscript with contributions from all authors.

## Acknowledgements

We thank Talia Karasov for advice in implementing the library preparation protocol. We greatly valued Derek Lundberg’s assistance with the liquid handling robot, which Frank Chan kindly let us use for our experiments. Hans de Jong offered helpful discussions regarding inversions. We thank all those who provided us with additional data (see Materials and Methods). This work was supported by the Max Planck Society.

## Supplemental Figure Legends

**Figure S1. Workflow diagram for genomic DNA library preparation using Nextera LITE.** Gel verification step is optional, but recommended. Workflow as presented requires about 2 - 3 hours of time. Multiple plates can be processed at the same time. Before sequencing, determine which plates have complementary sets of indices (see File S1) and can therefore be pooled into a single lane. Mix each individual 96-sample pool at an equimolar ratio in the final tube containing all libraries to be sequenced in a single lane.

**Figure S2. Read coverage distributions among samples.** A) Coverage plots for each 96-sample pool for all of the 1,920 Col X Ler F_2_ individuals produced for this study. B) Coverage plots for publicly available read data for additional F_2_ individuals derived from the same parental backgrounds categorized by their accession numbers at ArrayExpress. Data points are displayed in 0.2x coverage bins. Cross bar indicates the mean.

**Figure S3. CO resolution.** The interval window for CO breakpoint estimation (calculated by the distance between flanking markers determined by TIGER). Shown are the intervals for 16,709 of 17,077 total COs, with intervals > 20 kb omitted for ease of visualization.

**Figure S4. CO per pair as a function of chromosome length.** The equation of the line is 5.14*10^−8^*x* + 0.355 (R_2_ = 0.96). The gray shaded area represents the 95% confidence interval.

**Figure S5. Positions of *cis* double crossovers along chromosomes.** The positions of the first and second COs for pairs of double COs are plotted along the chromosomes. The chromosome number is indicated in the gray boxes to the right of the plots.

**Figure S6. CO frequencies adjacent to a 170-kb inversion on chromosome 3.** CO frequencies are plotted in 50-kb bins around the inversion (light gray shaded region).

**Figure S7. Distance to nearest CO position from the borders of SVs as a function of the size of the variant region.** Linear regressions revealed no significant correlation between variant size and the distance to the nearest CO position for any of the variants shown in the plots.

**Figure S8. Crossover rates in the 50-, 100-, and 200-kb up- and downstream of structural variants.** Rates are compared to the genome-wide means for the indicated window sizes (“Genome Mean” in plot) for inversions, insertions, deletions, transpositions (intrachrom) and translocations (interchrom) and copy number variations (CNV).

**Figure S9. Mean CO rates in the 200-kb flanking SVs.** Linear regressions revealed a significant correlation between SV size and the mean flanking CO rates only for copy number SVs (p = 0.046 with an R^2^ value of 0.045).

**Figure S10. Relationships among structural variants (SVs), “CO deserts”, and DNA methylation at CG positions.** A) Overlaps between CO-depleted regions “CO deserts” and SVs. Vertical black line indicates mean number of overlaps expected in 5000 random permutations. Vertical red line indicates the number of overlaps where p = 0.05. Vertical green line indicates the observed number of overlaps. Double-headed arrow highlights the difference between the mean of 5000 permutations and the observed number. B) Box plots showing the percentage of DNA methylation at CG positions in 10-kb windows that overlap with “CO deserts” (Desert) and those that do not (Outside) either with centromeric regions included or excluded. C) Box plots showing the percentage of DNA methylation at CG positions in 10-kb windows that overlap with structural variants (SV) and those that do not (Outside) either with centromeric regions included or excluded.

**Figure S11. Permutation tests of the overlaps between COs and NLR genes.** Permutation tests for all NLR loci (A) and subsets where the locus structure is the same between Col and Ler (B) and different between Col and Ler (C). For all plots, the vertical black line indicates mean number of overlaps expected in 5,000 random permutations. Vertical red line indicates the number of overlaps where p = 0.05. Vertical green line indicates the observed number of overlaps. Double arrow highlights the difference between the mean from 5,000 permutations and the observed number.

**Figure S12. COs in NLR genes relative to the number of copies in the locus.** Boxplots with individual data points display the CO rates within each locus type for loci where Col and Ler have the same number of copies in the locus (A) and loci where Col and Ler have different copy numbers (B).

**Figure S13. Locations of COs in six focal NLR loci.** These loci were also studied in (Choi *et al*. 2016).

**Figure S14. COs relative to ATAC-seq sites.** A) Permutation test of overlaps between COs and ATAC-seq sites. The vertical black line indicates mean number of overlaps expected in 1,000 random permutations. Vertical red line indicates the number of overlaps where p = 0.05. Vertical green line indicates the observed number of overlaps. Double-headed arrow highlights the difference between the mean of 1,000 permutations and the observed number. B) Distribution of distances from CO breakpoints to the nearest DNase HS site border. Vertical dashed green line indicates the median. The top 5% of distances are omitted from this plot for ease of visualization.

**Figure S15. Crossovers (COs) associated with sequence features.** A) CO rates (black) in comparison with Col/Ler SNP counts (grey shading) in 100-kb windows along the chromosomes and disease resistance genes (red ticks). CO rates are tallied as the number of COs per window divided by the total number of individuals. Horizontal dashed grey line indicates the genome-wide mean CO rate. B) Percentage of AT (red) and GC (blue) content across the genome. The green shading shows the CO rates presented in (A) and the same axis scale applies. Black ticks show disease resistance genes. The horizontal dashed red line indicates the genome-wide mean %AT content and the horizontal dashed blue line indicates the genome-wide mean %GC content. The position information shown in (B) also applies to (A). Solid vertical lines indicate chromosome boundaries and dashed vertical lines represent the mid-points of the centromeres in both A and B.

**File S1:** Oligos used for Nextera Indexing. All P7 oligos and P5 oligos S501-S508 contain standard Illumina index sequences. P5 oligos S509-S524 are from (Baym *et al*. 2015).

**File S2:** Flanking marker positions and CO positions. In the header, “chr” refers to the chromosome number, “block1.end” indicates the last marker of a genotype block, “block2.start” indicates the first marker of a new genotype block, “co.position” is the mid-point between the two markers flanking the CO. Of the 373 individuals that had been pre-selected by screening for lack of COs in the *420* reporter interval, 20% of individuals still had a CO in the region. Since this fraction was not substantially reduced compared with the fraction of non-selected individuals that had a CO in *420* (26%), we did not exclude them from our analysis. They are indicated in the column “select.420”.

**File S3:** List of genes in the top 95% longest CO deserts.

**File S4:** Comparison of the structures of NLR and other defense gene loci between Col and Ler.

## References

Alcázar R., M. von Reth, J. Bautor, E. Chae, D. Weigel, et al., 2014 Analysis of a plant complex resistance gene locus underlying immune-related hybrid incompatibility and its occurrence in nature. PLoS Genet. 10: e1004848.

Alhajturki D., S. Muralidharan, M. Nurmi, B. A. Rowan, J. E. Lunn, et al., 2018 Dose-dependent interactions between two loci trigger altered shoot growth in BG-5 × Krotzenburg-0 (Kro-0) hybrids of Arabidopsis thaliana. New Phytol. 217: 392–406.

Arabidopsis Genome Initiative, 2000 Analysis of the genome sequence of the flowering plant Arabidopsis thaliana. Nature 408: 796–815.

Bailey T. L., M. Boden, F. A. Buske, M. Frith, C. E. Grant, et al., 2009 MEME SUITE: tools for motif discovery and searching. Nucleic Acids Res. 37: W202–8.

Barton N. H., and B. Charlesworth, 1998 Why sex and recombination? Science 281: 1986–1990.

Baym M., S. Kryazhimskiy, T. D. Lieberman, H. Chung, M. M. Desai, et al., 2015 Inexpensive multiplexed library preparation for megabase-sized genomes. PLoS One 10: e0128036.

Blat Y., R. U. Protacio, N. Hunter, and N. Kleckner, 2002 Physical and Functional Interactions among Basic Chromosome Organizational Features Govern Early Steps of Meiotic Chiasma Formation. Cell 111: 791–802.

Bogdanov Y. F., T. M. Grishaeva, and S. Y. Dadashev, 2007 Similarity of the Domain Structure of Proteins as a Basis for the Conservation of Meiosis1, pp. 83–142 in International Review of Cytology, Academic Press.

Bolger A. M., M. Lohse, and B. Usadel, 2014 Trimmomatic: a flexible trimmer for Illumina sequence data. Bioinformatics 30: 2114–2120.

Burt A., 2000 Perspective: sex, recombination, and the efficacy of selection--was Weismann right? Evolution 54: 337–351.

Cao J., K. Schneeberger, S. Ossowski, T. Günther, S. Bender, et al., 2011 Whole-genome sequencing of multiple Arabidopsis thaliana populations. Nat. Genet. 43: 956–963.

Chae E., K. Bomblies, S.-T. Kim, D. Karelina, M. Zaidem, et al., 2014 Species-wide genetic incompatibility analysis identifies immune genes as hot spots of deleterious epistasis. Cell 159: 1341–1351.

Chakraborty U., and E. Alani, 2016 Understanding how mismatch repair proteins participate in the repair/anti-recombination decision. FEMS Yeast Res. 16: fow071.

Charlesworth D., 2016 The status of supergenes in the 21st century: recombination suppression in Batesian mimicry and sex chromosomes and other complex adaptations. Evol. Appl. 9: 74–90.

Chin D. B., R. Arroyo-Garcia, O. E. Ochoa, R. V. Kesseli, D. O. Lavelle, et al., 2001 Recombination and spontaneous mutation at the major cluster of resistance genes in lettuce (Lactuca sativa). Genetics 157: 831–849.

Choi K., X. Zhao, K. A. Kelly, O. Venn, J. D. Higgins, et al., 2013 Arabidopsis meiotic crossover hot spots overlap with H2A.Z nucleosomes at gene promoters. Nat. Genet. 45: 1327–1336.

Choi K., C. Reinhard, H. Serra, P. A. Ziolkowski, C. J. Underwood, et al., 2016 Recombination Rate Heterogeneity within Arabidopsis Disease Resistance Genes. PLoS Genet. 12: e1006179.

Choi K., X. Zhao, A. J. Tock, and C. Lambing, 2018 Nucleosomes and DNA methylation shape meiotic DSB frequency in Arabidopsis thaliana transposons and gene regulatory regions. Genome Research 28: 532–546.

Collins N., J. Drake, M. Ayliffe, Q. Sun, J. Ellis, et al., 1999 Molecular characterization of the maize Rp1-D rust resistance haplotype and its mutants. Plant Cell 11: 1365–1376.

Crown K. N., D. E. Miller, J. Sekelsky, and R. S. Hawley, 2018 Local Inversion Heterozygosity Alters Recombination throughout the Genome. Curr. Biol. 28: 2984–2990.

Dobzhansky T., 1931 The Decrease of Crossing-Over Observed in Translocations, and Its Probable Explanation. Am. Nat. 65: 214–232.

Ewels P., M. Magnusson, S. Lundin, and M. Käller, 2016 MultiQC: summarize analysis results for multiple tools and samples in a single report. Bioinformatics 32: 3047–3048.

Fang Z., T. Pyhäjärvi, A. L. Weber, R. K. Dawe, J. C. Glaubitz, et al., 2012 Megabase-scale inversion polymorphism in the wild ancestor of maize. Genetics 191: 883–894.

Fernandes J. B., M. Séguéla-Arnaud, C. Larchevêque, A. H. Lloyd, and R. Mercier, 2018 Unleashing meiotic crossovers in hybrid plants. Proc. Natl. Acad. Sci. U. S. A. 115: 2431–2436.

Fisher R. A., 1999 The Genetical Theory of Natural Selection: A Complete Variorum Edition. OUP Oxford.

Francis K. E., S. Y. Lam, B. D. Harrison, A. L. Bey, L. E. Berchowitz, et al., 2007 Pollen tetrad-based visual assay for meiotic recombination in Arabidopsis. Proc. Natl. Acad. Sci. U. S. A. 104: 3913–3918.

Fransz P. F., S. Armstrong, J. H. de Jong, L. D. Parnell, C. van Drunen, et al., 2000 Integrated cytogenetic map of chromosome arm 4S of A. thaliana: structural organization of heterochromatic knob and centromere region. Cell 100: 367–376.

Fransz P., G. Linc, C.-R. Lee, S. A. Aflitos, J. R. Lasky, et al., 2016 Molecular, genetic and evolutionary analysis of a paracentric inversion in Arabidopsis thaliana. Plant J. 88: 159–178.

Gel B., A. Díez-Villanueva, E. Serra, M. Buschbeck, M. A. Peinado, et al., 2016 regioneR: an R/Bioconductor package for the association analysis of genomic regions based on permutation tests. Bioinformatics 32: 289–291.

George C. M., and E. Alani, 2012 Multiple cellular mechanisms prevent chromosomal rearrangements involving repetitive DNA. Crit. Rev. Biochem. Mol. Biol. 47: 297–313.

Giraut L., M. Falque, J. Drouaud, L. Pereira, O. C. Martin, et al., 2011 Genome-wide crossover distribution in Arabidopsis thaliana meiosis reveals sex-specific patterns along chromosomes. PLoS Genet. 7: e1002354.

Gray J. C., and M. R. Goddard, 2012 Sex enhances adaptation by unlinking beneficial from detrimental mutations in experimental yeast populations. BMC Evol. Biol. 12: 43.

Hall J. C., 1972 Chromosome segregation influenced by two alleles of the meiotic mutant c(3)G in Drosophila melanogaster. Genetics 71: 367–400.

Hammarlund M., M. W. Davis, H. Nguyen, D. Dayton, and E. M. Jorgensen, 2005 Heterozygous insertions alter crossover distribution but allow crossover interference in Caenorhabditis elegans. Genetics 171: 1047–1056.

Hennig B. P., L. Velten, I. Racke, C. S. Tu, M. Thoms, et al., 2018 Large-Scale Low-Cost NGS Library Preparation Using a Robust Tn5 Purification and Tagmentation Protocol. G3 8: 79–89.

Herickhoff L., S. Stack, and J. Sherman, 1993 The relationship between synapsis, recombination nodules and chiasmata in tomato translocation heterozygotes. Heredity 71: 373–385.

Higgins J. D., M. Ferdous, K. Osman, and F. C. H. Franklin, 2011 The RecQ helicase AtRECQ4A is required to remove inter-chromosomal telomeric connections that arise during meiotic recombination in Arabidopsis. Plant J. 65: 492–502.

Hsu S. J., R. P. Erickson, J. Zhang, W. S. Garver, and R. A. Heidenreich, 2000 Fine linkage and physical mapping suggests cross-over suppression with a retroposon insertion at the npc1 mutation. Mamm. Genome 11: 774–778.

Ji X., 2014 Numerical and structural chromosome aberrations in cauliflower (Brassica oleracea var. botrytis) and Arabidopsis thaliana, (H. de Jong, Ed.)

Joron M., L. Frezal, R. T. Jones, N. L. Chamberlain, S. F. Lee, et al., 2011 Chromosomal rearrangements maintain a polymorphic supergene controlling butterfly mimicry. Nature 477: 203–206.

Keeney S., C. N. Giroux, and N. Kleckner, 1997 Meiosis-specific DNA double-strand breaks are catalyzed by Spo11, a member of a widely conserved protein family. Cell 88: 375–384.

Keeney S., 2001 Mechanism and control of meiotic recombination initiation. Curr. Top. Dev. Biol. 52: 1–53.

Kulathinal R. J., L. S. Stevison, and M. A. F. Noor, 2009 The genomics of speciation in Drosophila: diversity, divergence, and introgression estimated using low-coverage genome sequencing. PLoS Genet. 5: e1000550.

Küpper C., M. Stocks, J. E. Risse, N. Dos Remedios, L. L. Farrell, et al., 2016 A supergene determines highly divergent male reproductive morphs in the ruff. Nat. Genet. 48: 79–83.

Lamesch P., T. Z. Berardini, D. Li, D. Swarbreck, C. Wilks, et al., 2012 The Arabidopsis Information Resource (TAIR): improved gene annotation and new tools. Nucleic Acids Res. 40: D1202–10.

Li H., and R. Durbin, 2009 Fast and accurate short read alignment with Burrows-Wheeler transform. Bioinformatics 25: 1754–1760.

Li H., B. Handsaker, A. Wysoker, T. Fennell, J. Ruan, et al., 2009 The Sequence Alignment/Map format and SAMtools. Bioinformatics 25: 2078–2079.

Liu S., C.-T. Yeh, T. Ji, K. Ying, H. Wu, et al., 2009 Mu transposon insertion sites and meiotic recombination events co-localize with epigenetic marks for open chromatin across the maize genome. PLoS Genet. 5: e1000733.

Liu P., C. M. B. Carvalho, P. J. Hastings, and J. R. Lupski, 2012 Mechanisms for recurrent and complex human genomic rearrangements. Curr. Opin. Genet. Dev. 22: 211–220.

Long Q., F. A. Rabanal, D. Meng, C. D. Huber, A. Farlow, et al., 2013 Massive genomic variation and strong selection in Arabidopsis thaliana lines from Sweden. Nat. Genet. 45: 884–890.

Lowry D. B., and J. H. Willis, 2010 A widespread chromosomal inversion polymorphism contributes to a major life-history transition, local adaptation, and reproductive isolation. PLoS Biol. 8: e1000500.

Maher K. A., M. Bajic, K. Kajala, M. Reynoso, G. Pauluzzi, et al., 2018 Profiling of accessible chromatin regions across multiple plant species and cell types reveals common gene regulatory principles and new control modules. Plant Cell 30: 15–36.

Marand A. P., S. H. Jansky, H. Zhao, C. P. Leisner, X. Zhu, et al., 2017 Meiotic crossovers are associated with open chromatin and enriched with Stowaway transposons in potato. Genome Biol. 18: 203.

Massy B. de, 2013 Initiation of meiotic recombination: how and where? Conservation and specificities among eukaryotes. Annu. Rev. Genet. 47: 563–599.

Mather K., 1950 The Genetical Architecture of Heterostyly in Primula sinensis. Evolution 4: 340–352.

McClintock B., 1931 Cytological observations of deficiencies involving known genes, translocations and an inversion in Zea mays. University of Missouri, College of Agriculture, Agricultural Experiment Station.

McClintock B., 1933 The association of non-homologous parts of chromosomes in the mid-prophase of meiosis in zea mays. Z.Zellforsch 19: 191–237.

McDowell J. M., M. Dhandaydham, T. A. Long, M. G. Aarts, S. Goff, et al., 1998 Intragenic recombination and diversifying selection contribute to the evolution of downy mildew resistance at the RPP8 locus of Arabidopsis. Plant Cell 10: 1861–1874.

McKim K. S., A. M. Howell, and A. M. Rose, 1988 The effects of translocations on recombination frequency in Caenorhabditis elegans. Genetics 120: 987–1001.

Melamed-Bessudo C., E. Yehuda, A. R. Stuitje, and A. A. Levy, 2005 A new seed-based assay for meiotic recombination in Arabidopsis thaliana. Plant J. 43: 458–466.

Melamed-Bessudo C., S. Shilo, and A. A. Levy, 2016 Meiotic recombination and genome evolution in plants. Curr. Opin. Plant Biol. 30: 82–87.

Meyers B. C., A. Kozik, A. Griego, H. Kuang, and R. W. Michelmore, 2003 Genome-wide analysis of NBS-LRR-encoding genes in Arabidopsis. Plant Cell 15: 809–834.

Muller H. J., 1932 Some Genetic Aspects of Sex. Am. Nat. 66: 118–138.

Noël L., T. L. Moores, E. A. van Der Biezen, M. Parniske, M. J. Daniels, et al., 1999 Pronounced intraspecific haplotype divergence at the RPP5 complex disease resistance locus of Arabidopsis. Plant Cell 11: 2099–2112.

Oneal E., D. B. Lowry, K. M. Wright, Z. Zhu, and J. H. Willis, 2014 Divergent population structure and climate associations of a chromosomal inversion polymorphism across the Mimulus guttatus species complex. Mol. Ecol. 23: 2844–2860.

Parniske M., K. E. Hammond-Kosack, C. Golstein, C. M. Thomas, D. A. Jones, et al., 1997 Novel disease resistance specificities result from sequence exchange between tandemly repeated genes at the Cf-4/9 locus of tomato. Cell 91: 821–832.

Parniske M., and J. D. Jones, 1999 Recombination between diverged clusters of the tomato Cf-9 plant disease resistance gene family. Proc. Natl. Acad. Sci. U. S. A. 96: 5850–5855.

Peck J. R., 1994 A ruby in the rubbish: beneficial mutations, deleterious mutations and the evolution of sex. Genetics 137: 597–606.

Pisupati R., I. Reichardt, Ü. Seren, P. Korte, V. Nizhynska, et al., 2017 Verification of Arabidopsis stock collections using SNPmatch, a tool for genotyping high-plexed samples. Sci Data 4: 170184.

R Core Team, 2017 R: A Language and Environment for Statistical Computing

Richter T. E., T. J. Pryor, J. L. Bennetzen, and S. H. Hulbert, 1995 New rust resistance specificities associated with recombination in the Rp1 complex in maize. Genetics 141: 373–381.

Rodgers-Melnick E., P. J. Bradbury, R. J. Elshire, J. C. Glaubitz, C. B. Acharya, et al., 2015 Recombination in diverse maize is stable, predictable, and associated with genetic load. Proc. Natl. Acad. Sci. U. S. A. 112: 3823–3828.

Rowan B. A., V. Patel, D. Weigel, and K. Schneeberger, 2015 Rapid and inexpensive whole-genome genotyping-by-sequencing for crossover localization and fine-scale genetic mapping. G3 5: 385–398.

Rowan B. A., D. K. Seymour, E. Chae, D. S. Lundberg, and D. Weigel, 2017 Methods for Genotyping-by-Sequencing. Methods Mol. Biol. 1492: 221–242.

Salomé P. A., K. Bomblies, J. Fitz, R. A. E. Laitinen, N. Warthmann, et al., 2012 The recombination landscape in Arabidopsis thaliana F2 populations. Heredity 108: 447–455.

Schwander T., R. Libbrecht, and L. Keller, 2014 Supergenes and complex phenotypes. Curr. Biol. 24: R288–94.

Serra H., C. Lambing, C. H. Griffin, S. D. Topp, D. C. Nageswaran, et al., 2018 Massive crossover elevation via combination of HEI10 and recq4a recq4b during Arabidopsis meiosis. Proc. Natl. Acad. Sci. U. S. A. 115: 2437–2442.

Shilo S., C. Melamed-Bessudo, Y. Dorone, N. Barkai, and A. A. Levy, 2015 DNA Crossover Motifs Associated with Epigenetic Modifications Delineate Open Chromatin Regions in Arabidopsis. Plant Cell 27: 2427–2436.

Stevison L. S., K. B. Hoehn, and M. A. F. Noor, 2011 Effects of Inversions on Within- and Between-Species Recombination and Divergence. Genome Biol. Evol. 3: 830–841.

Sturtevant A. H., 1921 A Case of Rearrangement of Genes in Drosophila. Proc. Natl. Acad. Sci. U. S. A. 7: 235–237.

Sturtevant A. H., 1926 A crossover reducer in Drosophila melanogaster due to inversion of a section of the third chromosome. Biol. Zent. Bl. 46: 697–702.

Sullivan A. M., A. A. Arsovski, J. Lempe, K. L. Bubb, M. T. Weirauch, et al., 2014 Mapping and dynamics of regulatory DNA and transcription factor networks in A. thaliana. Cell Rep. 8: 2015–2030.

Sun Q., N. C. Collins, M. Ayliffe, S. M. Smith, J. Drake, et al., 2001 Recombination between paralogues at the Rp1 rust resistance locus in maize. Genetics 158: 423–438.

Sybenga J., 1970 Simultaneous negative and positive chiasma interference across the break point in interchange heterozygotes. Genetica 41: 209–230.

Szostak J. W., T. L. Orr-Weaver, R. J. Rothstein, and F. W. Stahl, 1983 The double-strand-break repair model for recombination. Cell 33: 25–35.

Thompson M. J., and C. D. Jiggins, 2014 Supergenes and their role in evolution. Heredity 113: 1–8.

Underwood C. J., K. Choi, C. Lambing, X. Zhao, H. Serra, et al., 2018 Epigenetic activation of meiotic recombination near Arabidopsis thaliana centromeres via loss of H3K9me2 and non-CG DNA methylation. Genome Res. 28: 519–531.

Van de Weyer A.-L., F. Monteiro, O. J. Furzer, M. T. Nishimura, V. Cevik, et al., 2019 The Arabidopsis thaliana pan-NLRome. bioRxiv 537001.

Wijnker E., G. Velikkakam James, J. Ding, F. Becker, J. R. Klasen, et al., 2013 The genomic landscape of meiotic crossovers and gene conversions in Arabidopsis thaliana. Elife 2: e01426.

Wulff B. B. H., C. M. Thomas, M. Parniske, and J. D. G. Jones, 2004 Genetic variation at the tomato Cf-4/Cf-9 locus induced by EMS mutagenesis and intralocus recombination. Genetics 167: 459–470.

Yang S., L. Wang, J. Huang, X. Zhang, Y. Yuan, et al., 2015 Parent–progeny sequencing indicates higher mutation rates in heterozygotes. Nature 523: 463–467.

Yelina N. E., K. Choi, L. Chelysheva, M. Macaulay, B. de Snoo, et al., 2012 Epigenetic remodeling of meiotic crossover frequency in Arabidopsis thaliana DNA methyltransferase mutants. PLoS Genet. 8: e1002844.

Yelina N. E., C. Lambing, T. J. Hardcastle, X. Zhao, B. Santos, et al., 2015 DNA methylation epigenetically silences crossover hot spots and controls chromosomal domains of meiotic recombination in Arabidopsis. Genes Dev. 29: 2183–2202.

Zapata L., J. Ding, E.-M. Willing, B. Hartwig, D. Bezdan, et al., 2016 Chromosome-level assembly of Arabidopsis thaliana Ler reveals the extent of translocation and inversion polymorphisms. Proc. Natl. Acad. Sci. U. S. A. 113: E4052–60.

Zickler D., and N. Kleckner, 1999 Meiotic chromosomes: integrating structure and function. Annu. Rev. Genet. 33: 603–754.

Ziolkowski P. A., L. E. Berchowitz, C. Lambing, N. E. Yelina, X. Zhao, et al., 2015 Juxtaposition of heterozygous and homozygous regions causes reciprocal crossover remodelling via interference during Arabidopsis meiosis. Elife 4: e03708.

