## Supplementary material for "An ultra high-density *Arabidopsis thaliana* crossover map that refines the influences of structural variation and epigenetic features": Tables S1-S6 and Figures S1-S15

This supplement includes:
Tables S1 – S6
Figures S1 - S15

**Table S1. Genetic distances.**

| Interval | Chr | Starting position | Ending position | cM (Whole genome) | cM (Reporters) |
|---|---|---|---|---|---|
| *I1b* | 1 | 3,905,441 | 5,755,618 | 4.9 | 6.6 |
| *I1fg* | 1 | 24,645,163 | 25,956,590 | 5.6 | 8.2 |
| *I2f* | 2 | 18,286,716 | 18,957,093 | 3.3 | 7.6 |
| *420* | 3 | 256,516 | 5,361,637 | 14.3 | 12.3 |
| *CEN3* | 3 | 11,115,724 | 16,520,560 | 10.2 | 11.9 |

Comparison of genetic distances in reporter intervals estimated from the whole-genome analysis presented in this study compared with those previously estimated from fluorescent reporters (Ziolkowski et al. 2015). Note that the estimates for the intervals *I1b*, *I1fg*, *I2f*, and *CEN3* are based on pollen reporters and represent male meiosis only. The whole genome data calculated from the study of crossovers (COs) in $F_2$ plants and the *420* interval in both datasets reflect sex-averaged genetic distances.

**Table S2. Overlaps between CO intervals and structural variants**

| Variant type | Total Number | Number of overlaps with CO intervals (observed) | Mean number of overlaps with CO intervals (permuted) |
|---|---:|---:|---:|
| Inversion | 38 | 13* | 246.4 |
| Insertion | 311 | 134* | 231.8 |
| Deletion | 426 | 156* | 238 |
| Intrachromosomal Translocations | 102 | 113* | 255.6 |
| Interchromosomal Translocations | 271 | 159* | 297.8 |
| Copy Number Variations | 67 | 38 | 53.8 |

*p < 0.05 after 5000 permutations

**Table S3. Crossover rates in flanking regions up- and downstream of structural variants**

| | 50 kb CO rate (cM/Mb) | 100 kb CO rate (cM/Mb) | 200 kb CO rate (cM/Mb) |
|---|---|---|---|
| Genome mean | 3.2 | 3.2 | 3.2 |
| Inversion | 3.4 | 3.6 | 3.8 |
| Insertion | 4.3 | 4.3 | 4.1 |
| Deletion | 4.3 | 4.2 | 4.1 |
| Intrachromosomal Translocation | 3.7 | 3.8 | 4.1 |
| Interchromosomal Translocation | 4.3 | 4.4 | 4.4 |
| Copy Number Variation | 4.4 | 4.2 | 4.0 |

**Table S4. GO Slim Enrichment in CO deserts**

| PANTHER GO-Slim Biological Process | Arabidopsis thaliana reference | CO Desert Set Observed | CO Desert Set Expected | Fold Enrichment | Raw P-value | FDR |
|---|---|---|---|---|---|---|
| meiosis (GO:0007126) | 60 | 15 | 4.86 | 3.08 | 3.93E-04 | 1.23E-02 |
| cellular component movement (GO:0006928) | 131 | 24 | 10.62 | 2.26 | 6.22E-04 | 1.67E-02 |
| DNA repair (GO:0006281) | 189 | 33 | 15.32 | 2.15 | 1.60E-04 | 6.01E-03 |
| DNA metabolic process (GO:0006259) | 351 | 57 | 28.45 | 2 | 6.39E-06 | 4.01E-04 |

**Table S5. CO frequency in defense genes**

| **Gene Annotation** | **Number of Genes** | **Number of Genes with COs** | **Proportion of Genes with COs** |
|---|---|---|---|
| CNL | 26 | 9 | 0.35 |
| NB-ARC | 15 | 6 | 0.4 |
| NB-LRR | 10 | 1 | 0.1 |
| P-loop protein | 6 | 0 | 0 |
| Phloem protein | 4 | 0 | 0 |
| TIR-NBS | 13 | 2 | 0.15 |
| TIR family | 20 | 5 | 0.25 |
| TNL | 87 | 29 | 0.33 |
| Transmembrane receptors | 4 | 1 | 0.25 |
| Other | 12 | 2 | 0.17 |

**Table S6. CO Frequency of sequence motifs associated with a random subset of 2500 crossovers**

| Motif | Number | Percent |
|---|---|---|
| PolyA/T | 2102 | 84.1 |
| CTT/GAA | 1141 | 45.6 |
| CT/GA | 943 | 37.0 |
| CCN/GGN | 304 | 12.2 |
| PolyA/T or CTT/GAA | 2292 | 91.7 |
| All four motifs combined | 2363 | 94.5 |

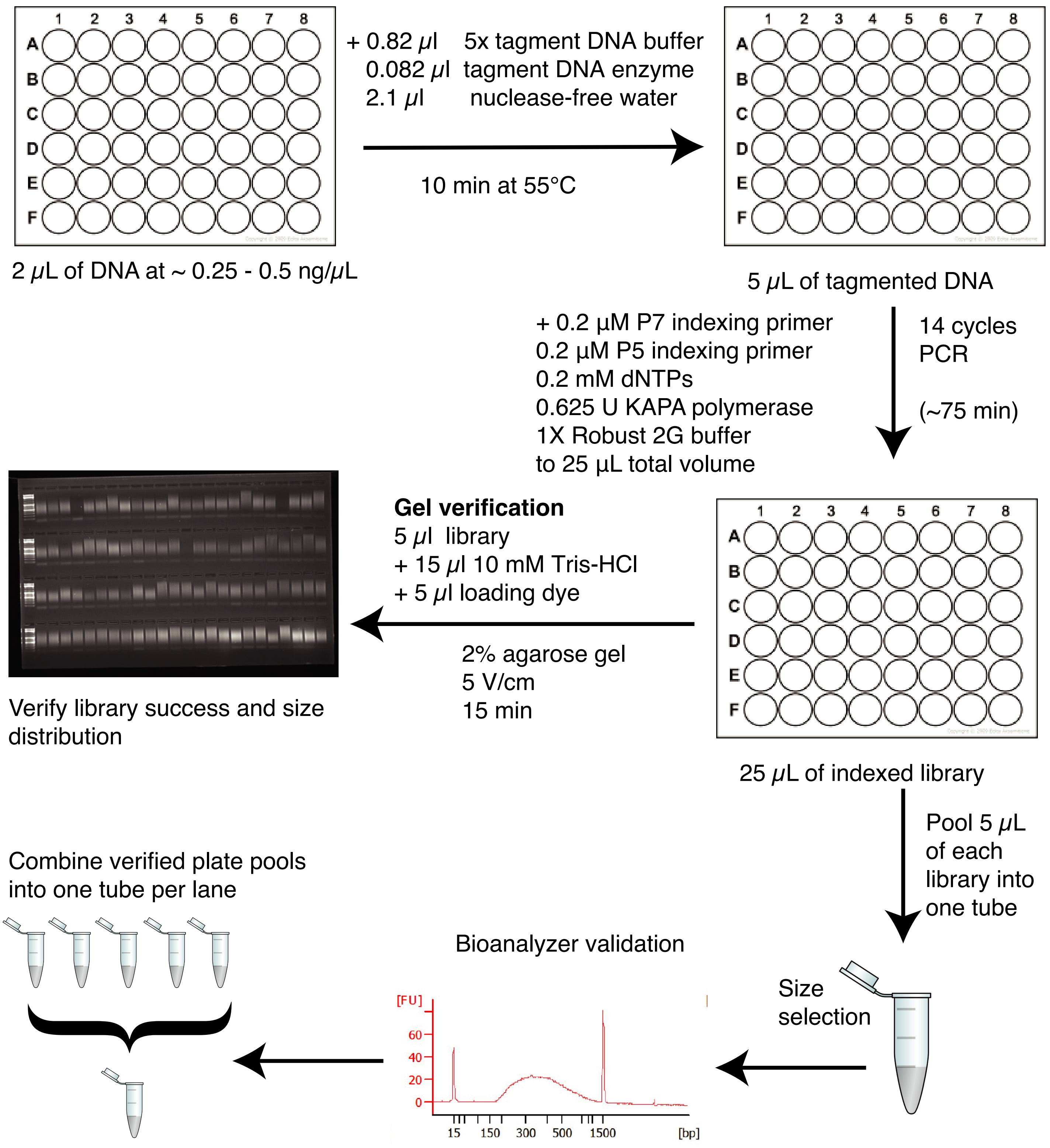


**Figure S1. Workflow diagram for genomic DNA library preparation using Nextera LITE.** Gel verification step is optional, but recommended. Workflow as presented requires about 2 - 3 hours of time. Multiple plates can be processed at the same time. Before sequencing, determine which plates have complementary sets of indices (see File S1) and can therefore be pooled into a single lane. Mix each individual 96-sample pool at an equimolar ratio in the final tube containing all libraries to be sequenced in a single lane.

**A**

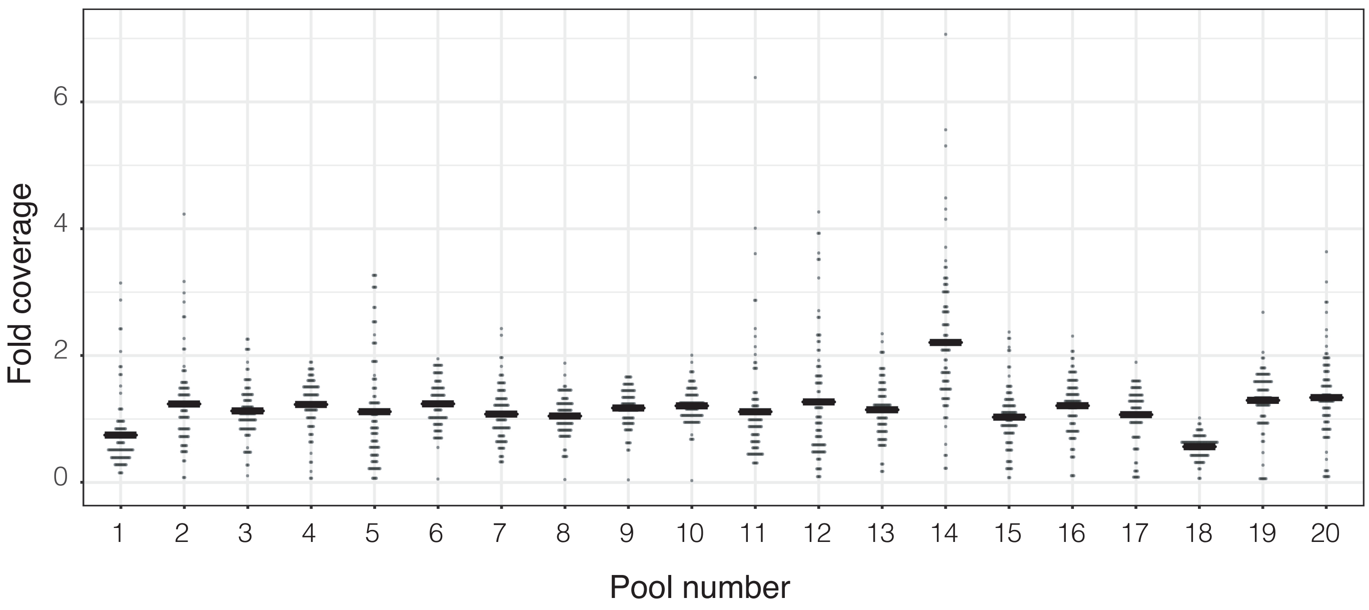


**B**

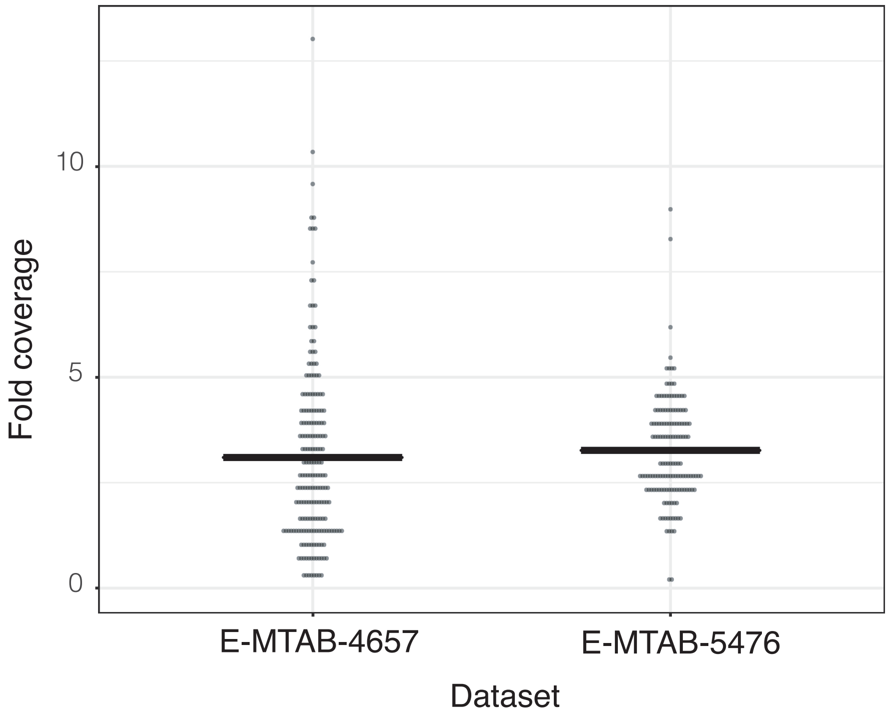


**Figure S2. Coverage distributions among samples**. A) Coverage plots for each 96-sample pool for all of the 1920 Col X Ler $F_2$ individuals produced for this study. B) Coverage plots for publicly available read data for additional $F_2$ individuals derived from the same parental backgrounds categorized by their accession numbers at ArrayExpress. Data points are displayed in 0.2x coverage bins. Cross bar indicates the mean.

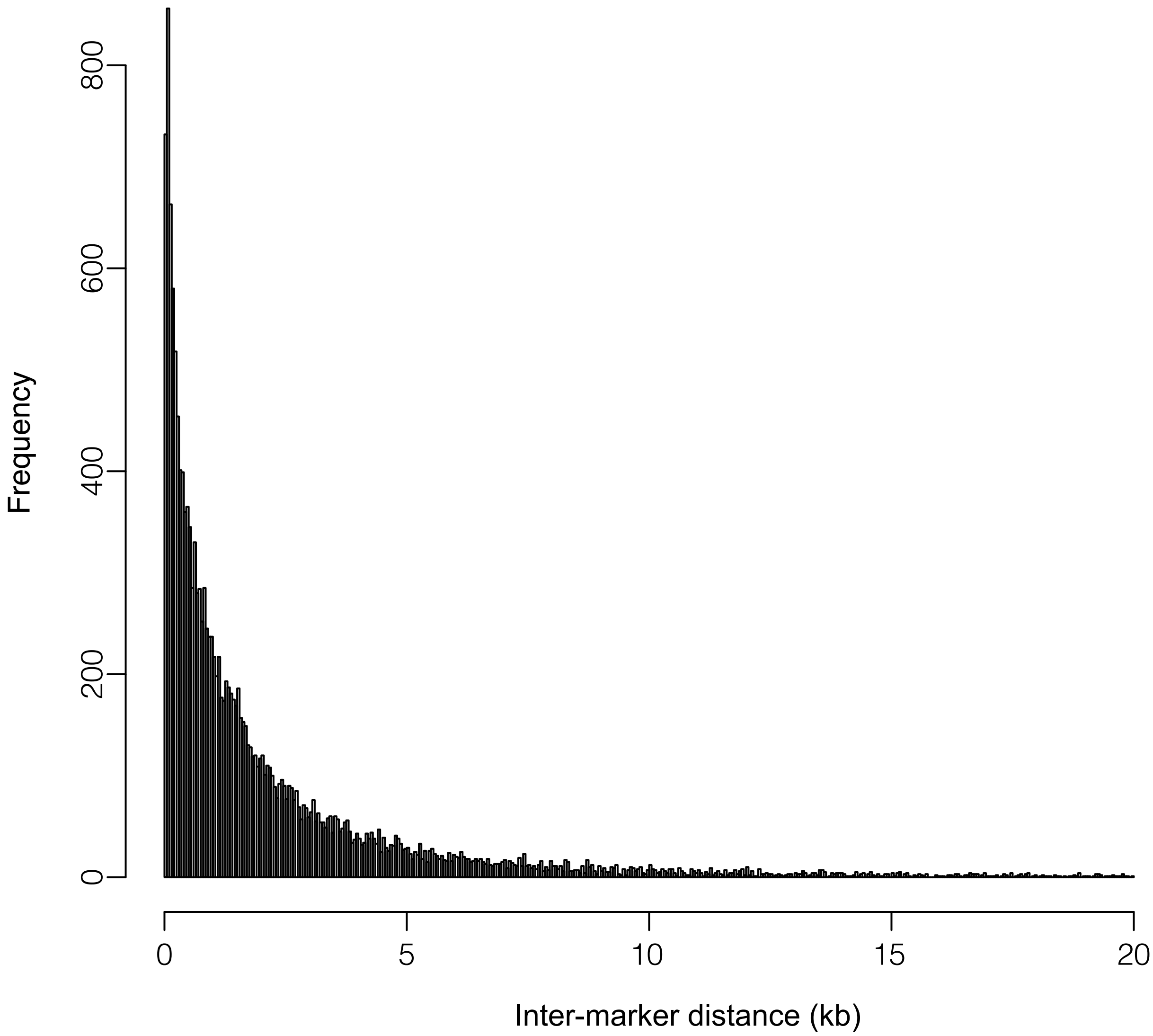


**Figure S3. Crossover resolution.** The interval window for CO breakpoint estimation (calculated by the distance between flanking markers determined by TIGER. Shown are the intervals for 16,709 of 17,077 total COs, as intervals > 20 kb have been omitted for ease of visualization.

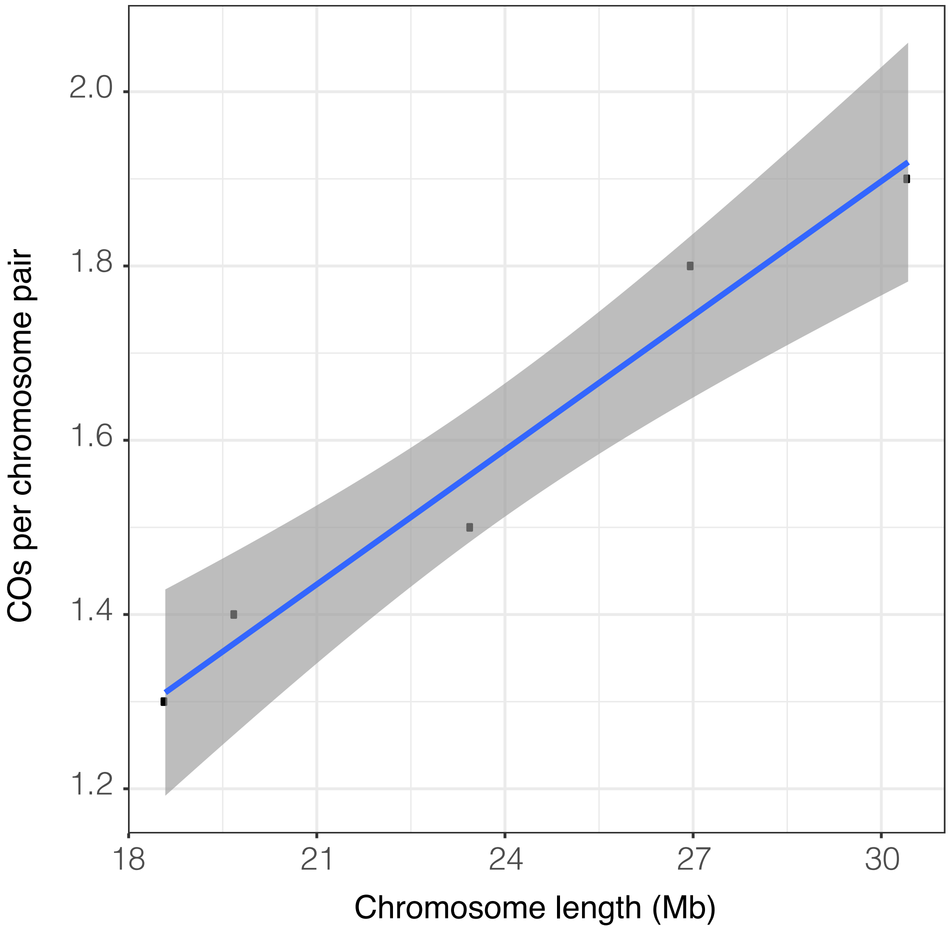


**Figure S4. Crossovers per pair as a function of chromosome length.** The equation of the line is 5.14*10$^{-08}$ + 3.55 ($R^2$ = 0.96). The gray shaded area represents the 95% confidence interval.

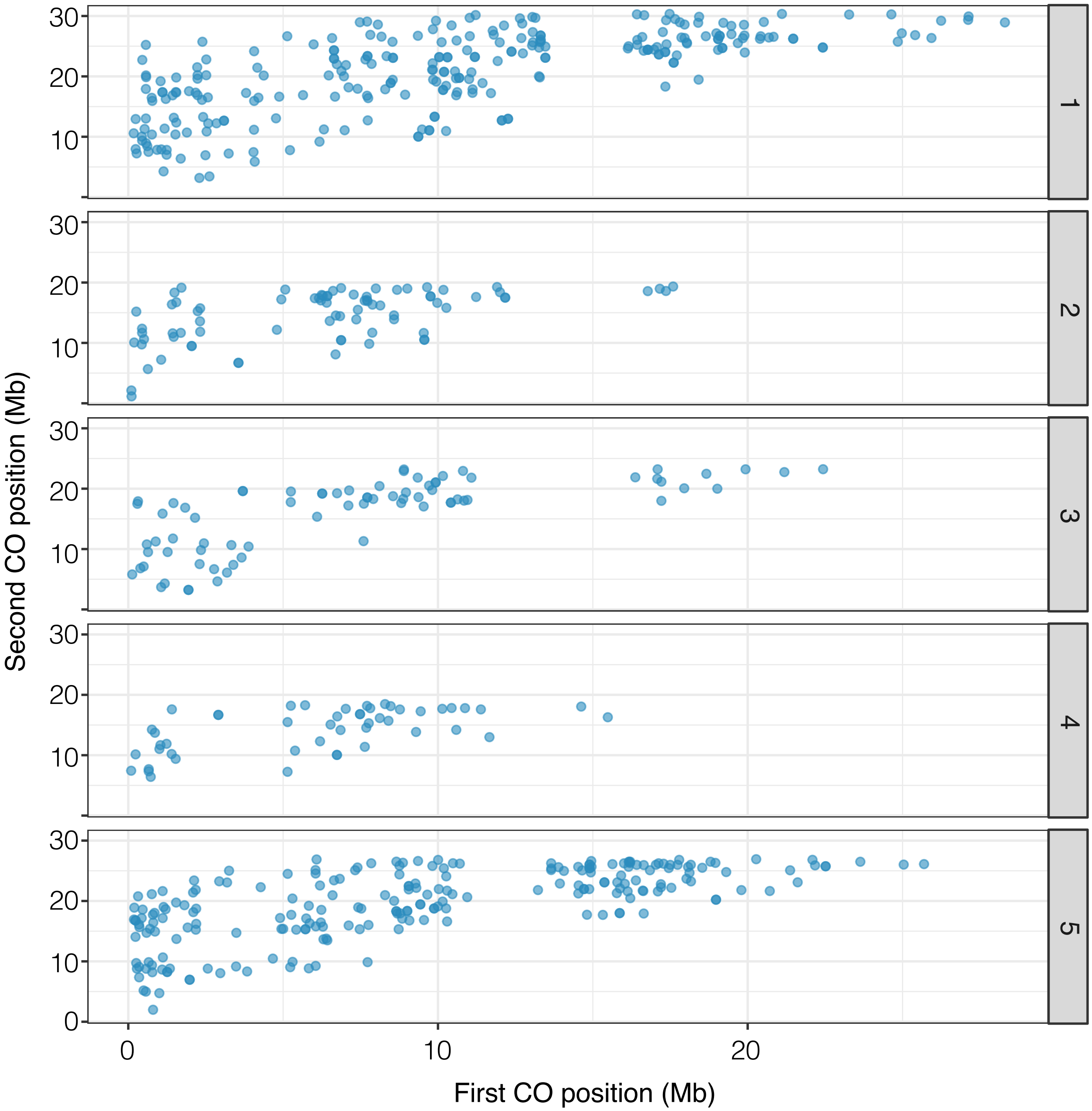


**Figure S5. Positions of cis double crossovers along chromosomes.** The positions of the first and second COs for pairs of double COs are plotted along the chromosomes. The chromosome number is indicated in the gray boxes to the right of the plots.

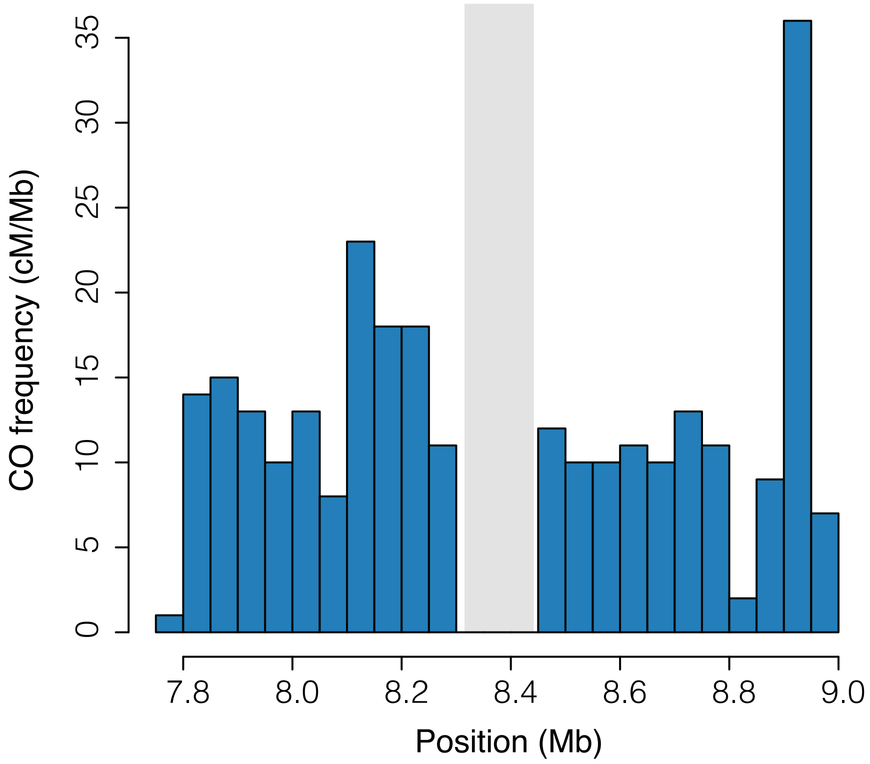


**Figure S6. Crossover frequencies adjacent to a 170-kb inversion on chromosome 3.** CO frequencies (cM/Mb) are plotted in 50-kb bins around the inversion (light gray shaded region).

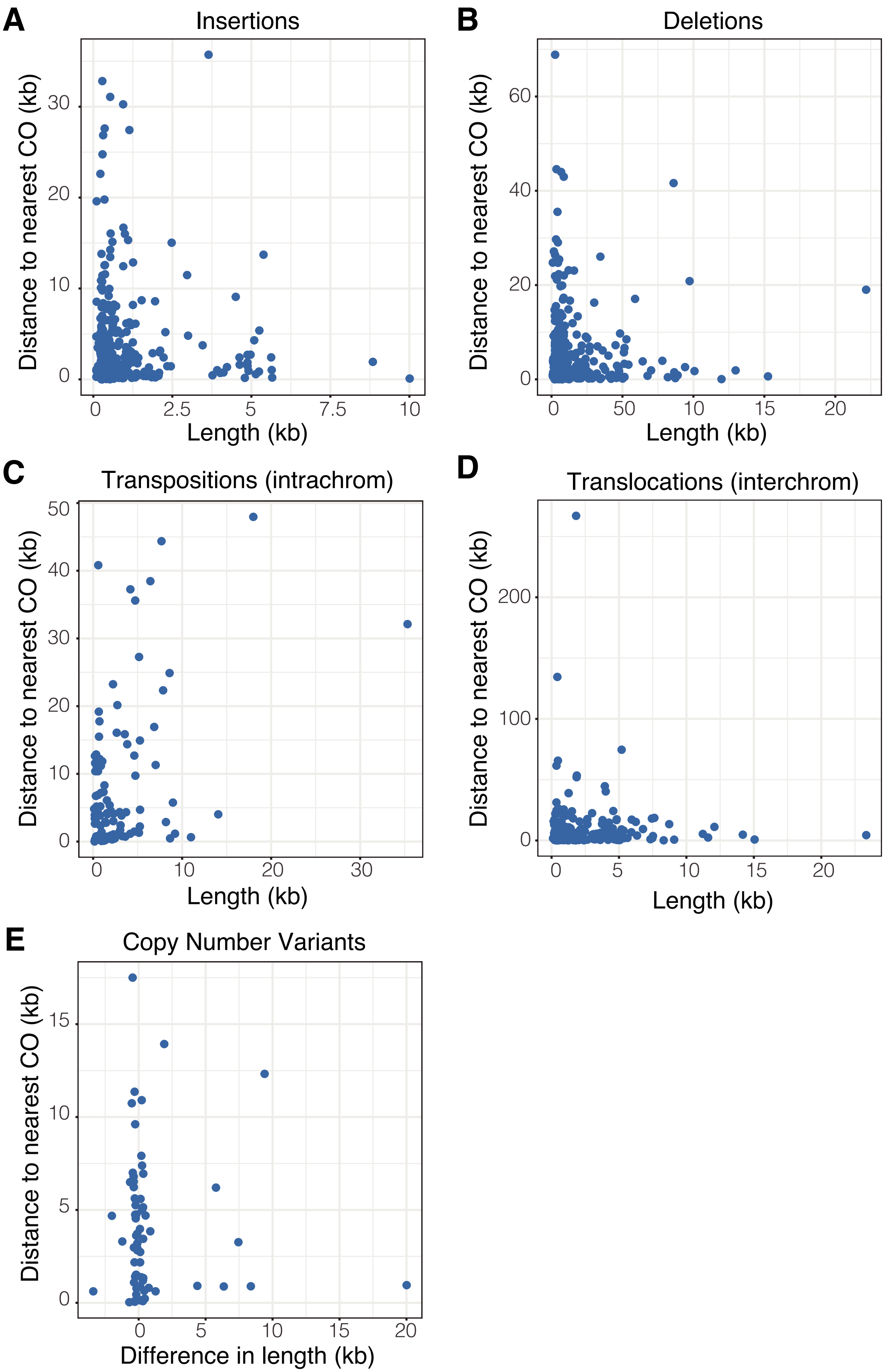


**Figure S7. Distance to nearest crossover position from the borders of structural variants as a function of the size of the variant region**. Linear regressions revealed no significant correlation between variant size and the distance to the nearest CO position for any of the variants shown in the plots.

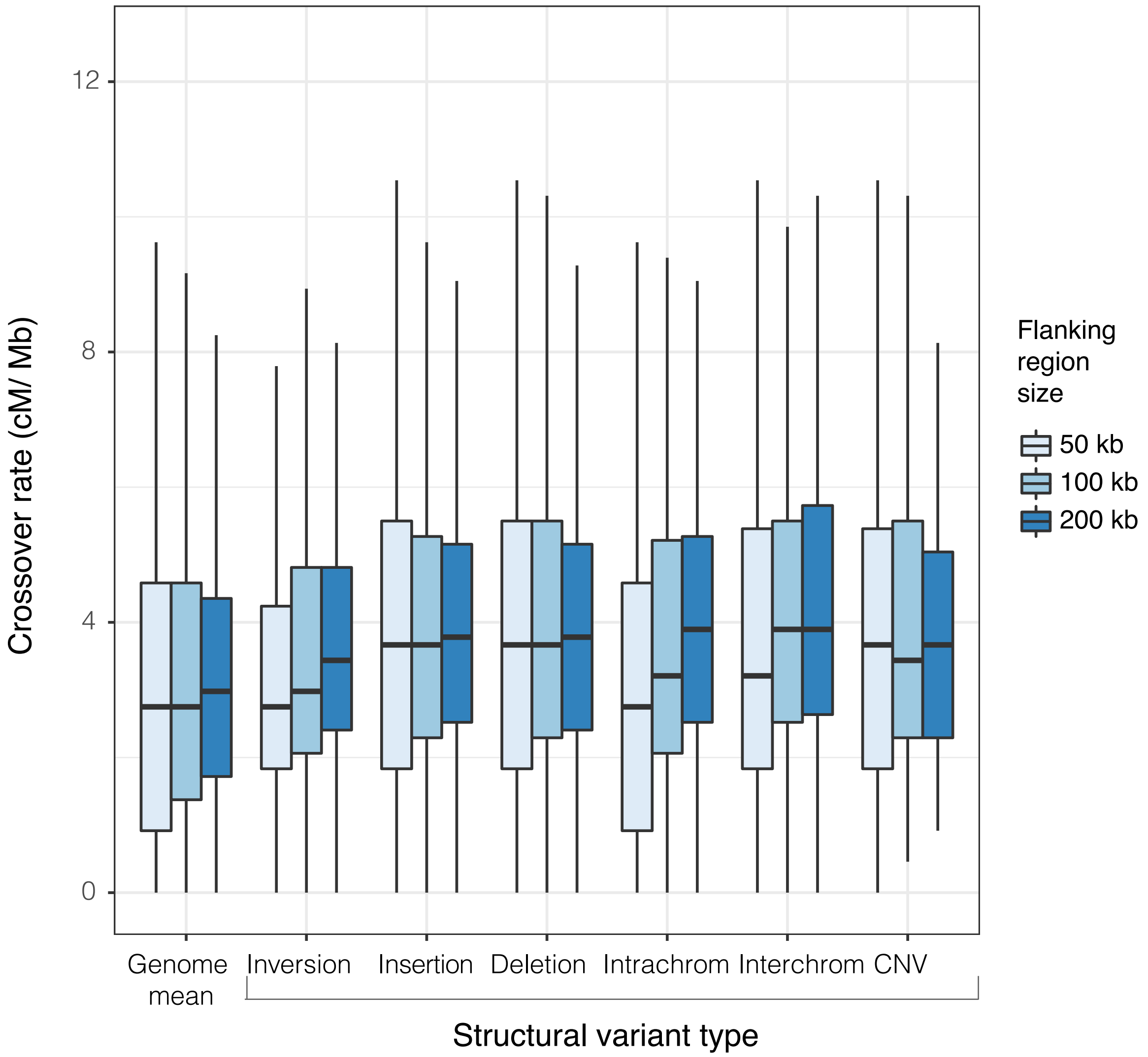


**Figure S8. Crossover rates in the 50-, 100-, and 200-kb up- and downstream of structural variants.** Rates are compared to the genome-wide means for the indicated window sizes ("Genome Mean" in plot) for inversions, insertions, deletions, transpositions (intrachrom) and translocations (interchrom) and copy number variations (CNV).

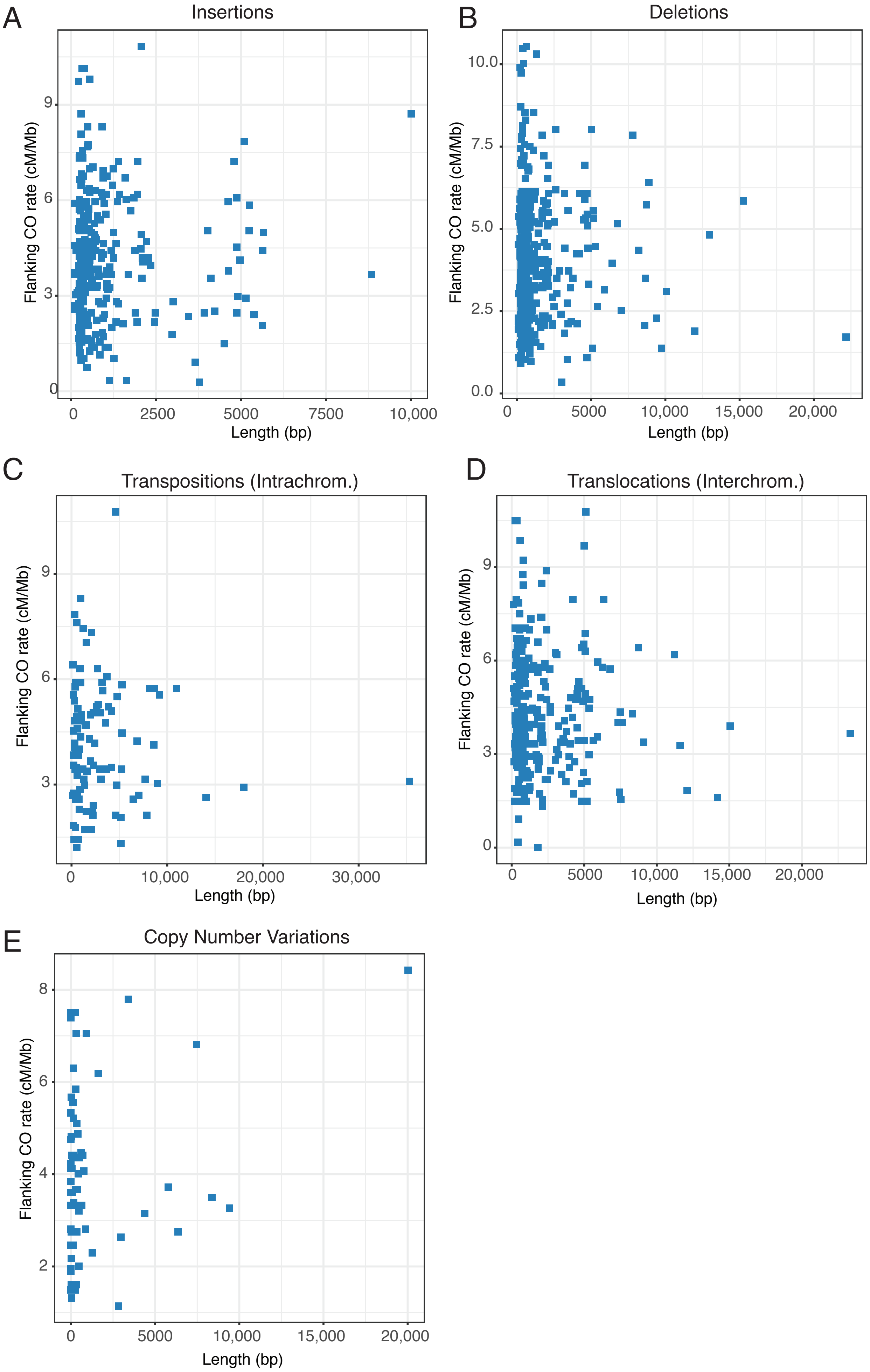


**Figure S9. Mean CO rates in the 200-kb up- and downstream of structural variants**. Linear regressions revealed a significant correlation between variant size and the mean flanking CO rates only for copy number variations ($p = 0.046$ with an $R^2$ value of 0.045).

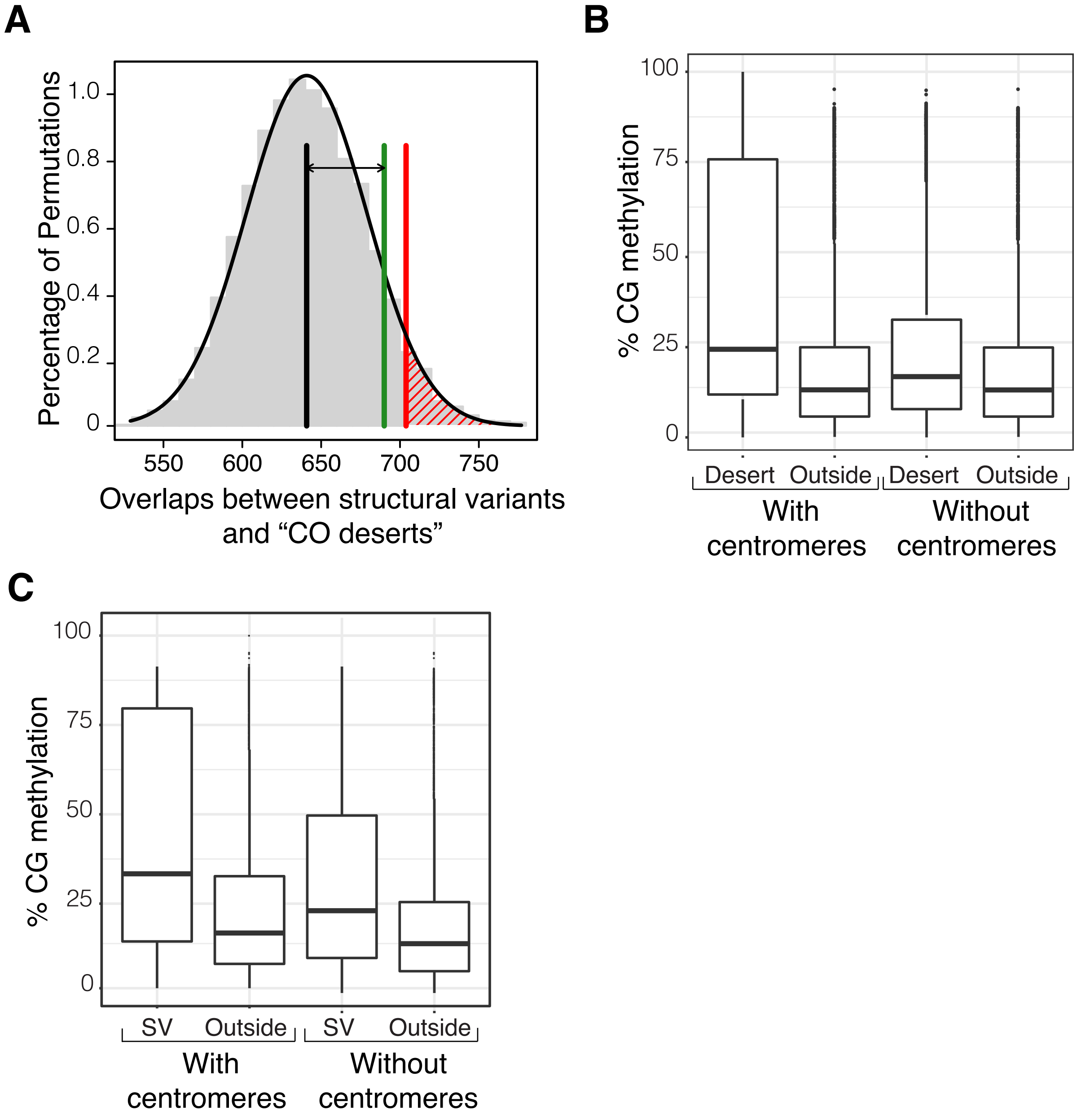


**Figure S10. Relationships among structural variants (SVs), "CO deserts", and DNA methylation at CG positions**. A) Overlaps between CO-depleted regions "CO deserts" and SVs. Vertical black line indicates mean number of overlaps expected in 5000 random permutations. Vertical red line indicates the number of overlaps where p = 0.05. Vertical green line indicates the observed number of overlaps. Double-headed arrow highlights the difference between the mean of 5000 permutations and the observed number. B) Box plots showing the percentage of DNA methylation at CG positions in 10-kb windows that overlap with "CO deserts" (Desert) and those that do not (Outside) either with centromeric regions included or excluded. C) Box plots showing the percentage of DNA methylation at CG positions in 10-kb windows that overlap with structural variants (SV) and those that do not (Outside) either with centromeric regions included or excluded.

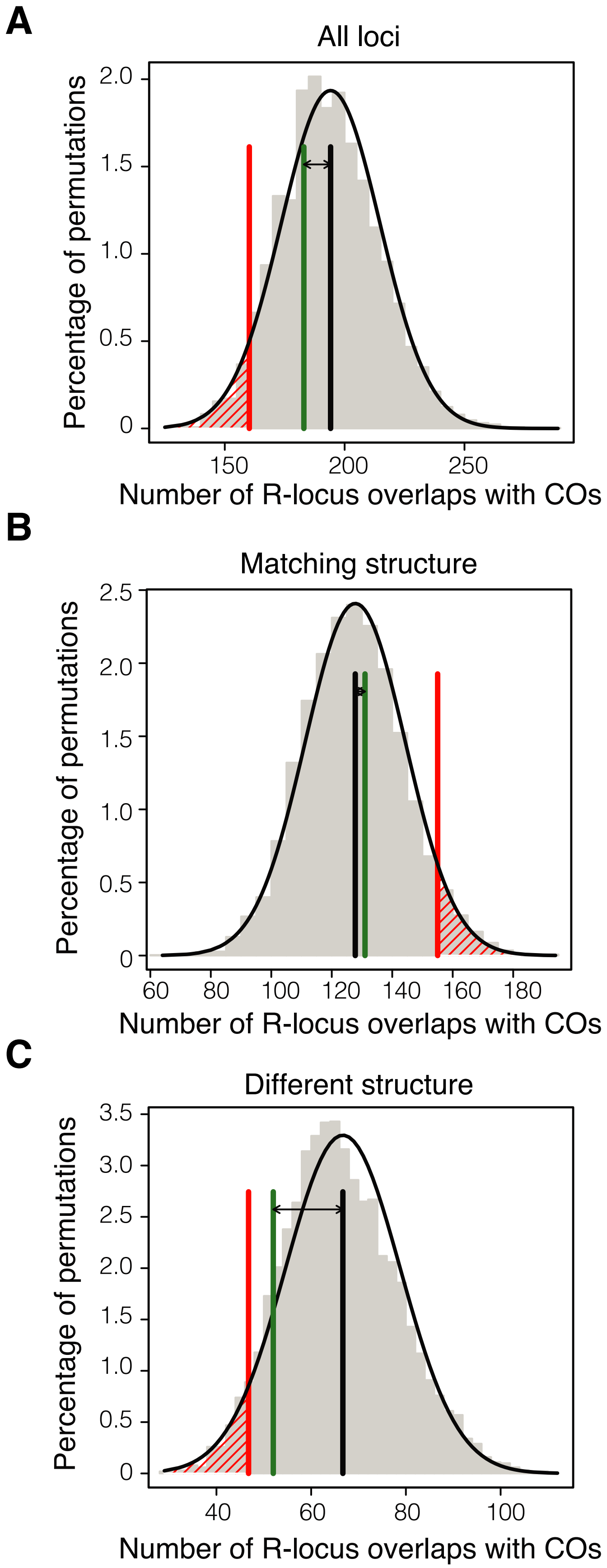


**Figure S11. Permutation tests of the overlaps between crossovers and R genes**. Permutation tests for all R-gene loci (A) and subsets where the locus structure is the same between Col and Ler (B) and different between Col and Ler (C). For all plots, the vertical black line indicates mean number of overlaps expected in 5000 random permutations. Vertical red line indicates the number of overlaps where p = 0.05. Vertical green line indicates the observed number of overlaps. Double-headed arrow highlights the difference between the mean of 5000 permutations and the observed number.

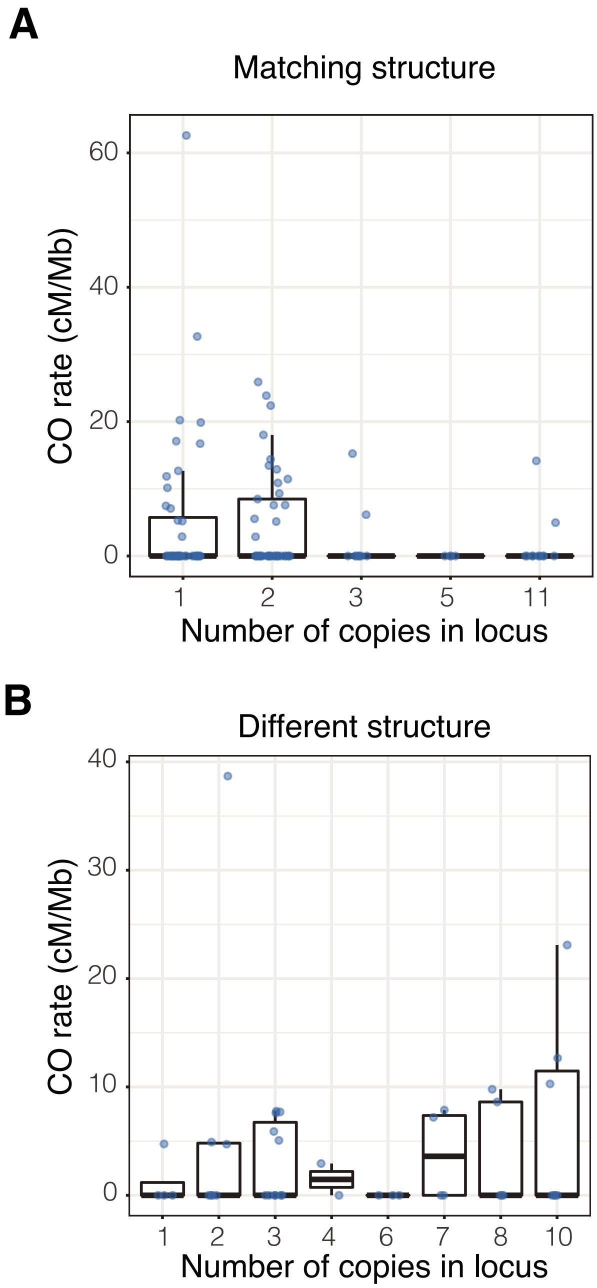


**Figure S12. Crossovers in R-genes relative to the number of copies in the the locus.** Boxplots with individual data points display the CO rates within each locus type for loci where Col and Ler have the same number of copies in the locus (A) and loci where Col and Ler have different numbers of copies in the locus (B).

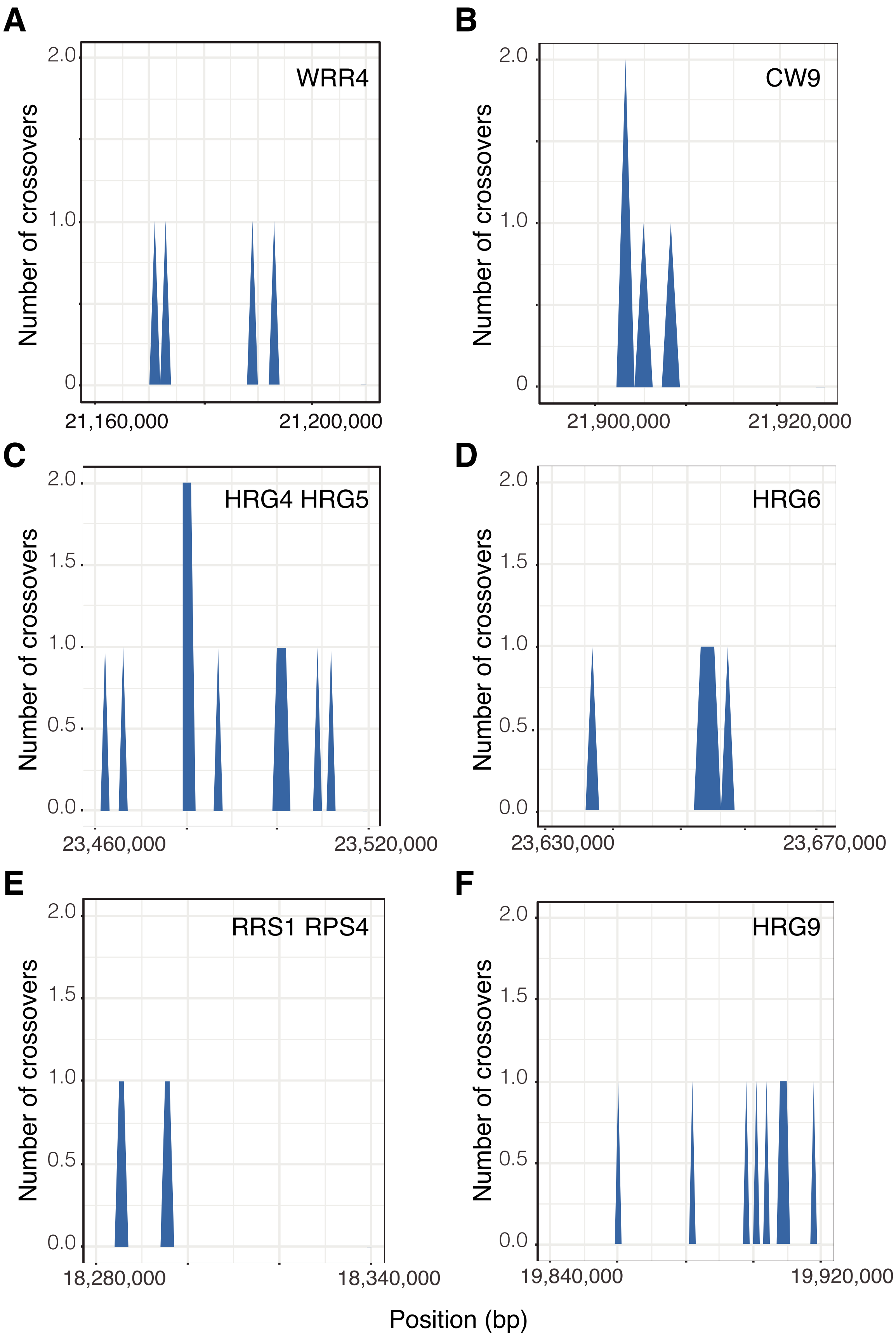


**Figure S13. Locations of crossovers in six focal R-gene loci.** R-gene loci were previously studied in Choi et al. (2016).

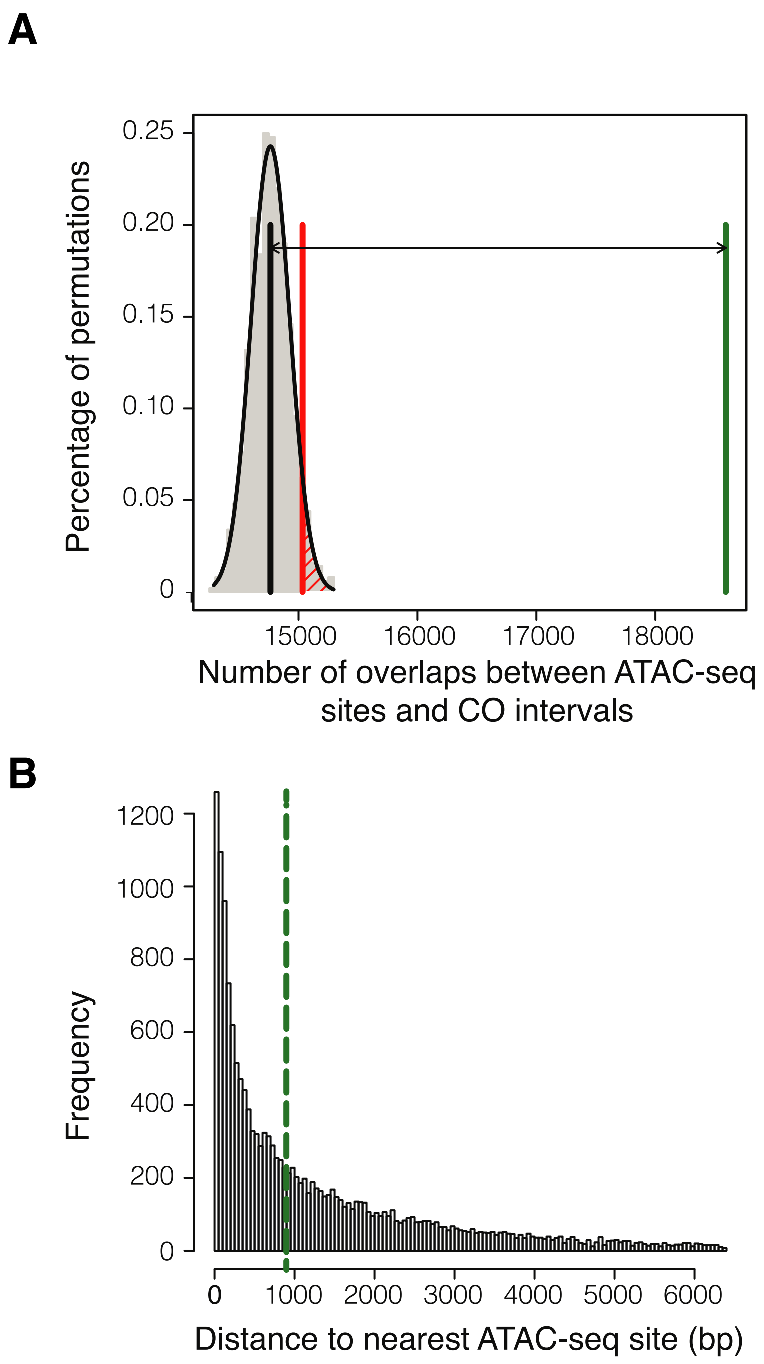


**Figure S14. Crossovers relative to ATAC-seq sites.** A) Permutation test of overlaps between COs and ATAC-seq sites. The vertical black line indicates mean number of overlaps expected in 1000 random permutations. Vertical red line indicates the number of overlaps where p = 0.05. Vertical green line indicates the observed number of overlaps. Double-headed arrow highlights the difference between the mean of 1000 permutations and the observed number. B) Distribution of distances from CO breakpoints to the nearest DNase HS site border. Vertical dashed green line indicates the median. The greatest 5% of distances are omitted from this plot for ease of visualization.

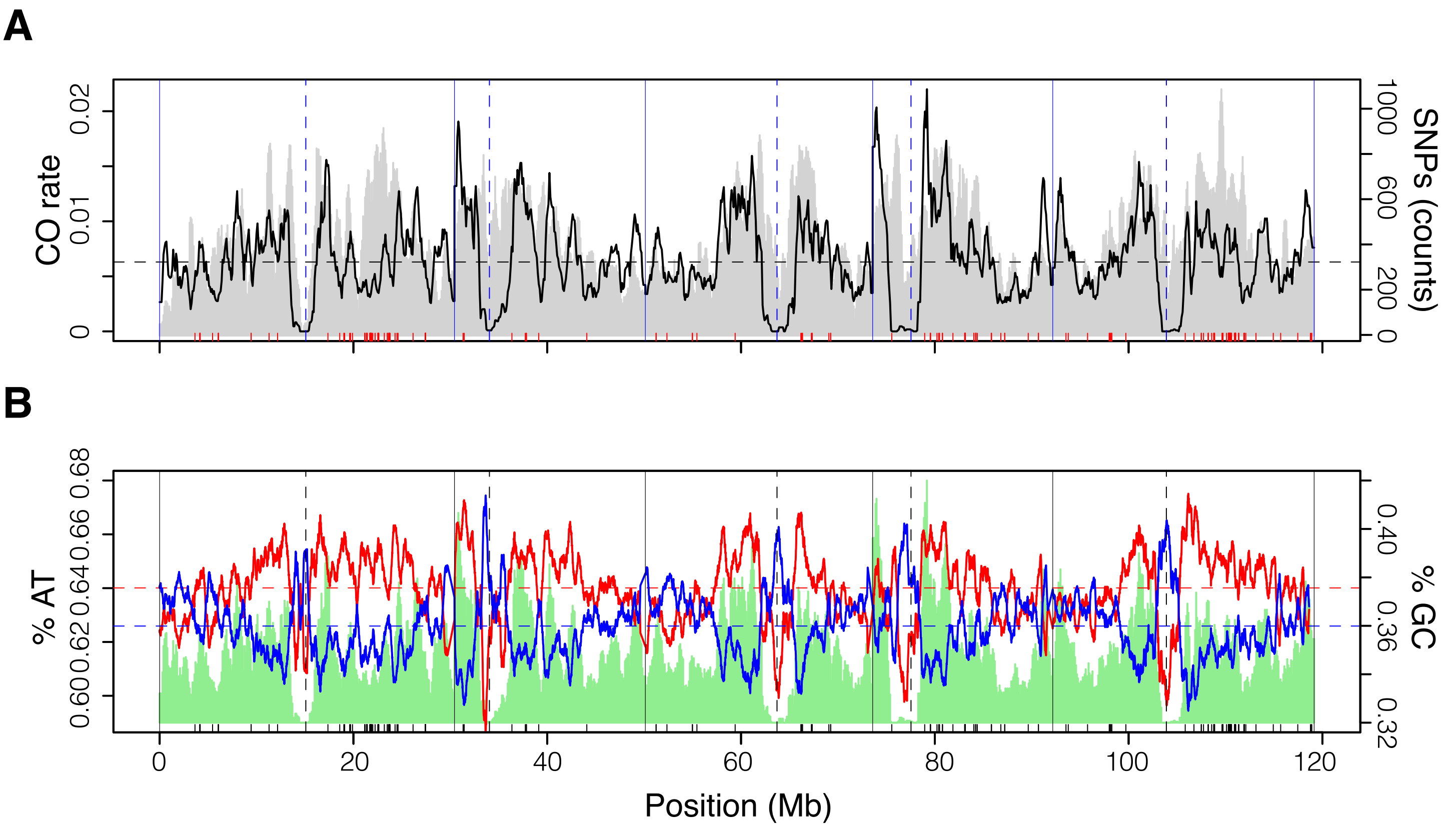


**Figure S15. Crossovers (COs) associated with sequence features**. A) CO rates (black) in comparison with Col/Ler SNP counts (grey shading) in 100-kb windows along the chromosomes and disease resistance genes (red ticks). CO rates are tallied as the number of COs per window divided by the total number of individuals. Horizontal dashed grey line indicates the genome-wide mean CO rate. B) Percentage of AT (red) and GC (blue) content across the genome. The green shading shows the CO rates presented in (A) and the same axis scale applies. Black ticks show disease resistance genes. The horizontal dashed red line indicates the genome-wide mean %AT content and the horizontal dashed blue line indicates the genome-wide mean %GC content. The position information shown in (B) also applies to (A). Solid vertical lines indicate chromosome boundaries and dashed vertical lines represent the mid-points of the centromeres in both A and B.
